# A single-cell transcriptomic atlas of the Echinococcus multilocularis metacestode reveals cellular diversity and molecular specialization

**DOI:** 10.64898/2026.08.23.746249

**Authors:** Julia A. Loos, Monika Bergmann, Arturo Calderón-Gallegos, Klaus Brehm

**Author notes:** Corresponding authors: Julia A. Loos: Consultant Laboratory for Echinococcosis, Institute of Hygiene and Microbiology, Josef-Schneider-Straße 2, 97070, Würzburg, Germany., Klaus Brehm: Consultant Laboratory for Echinococcosis, Institute of Hygiene and Microbiology, Josef-Schneider-Straße 2, 97070, Würzburg, Germany.

## Abstract

The metacestode of *Echinococcus multilocularis* is the proliferative larval stage responsible for alveolar echinococcosis and displays remarkable capacities for long-term growth, regeneration and development within the host. Despite its medical relevance, the cellular composition and molecular organization of this stage remain incompletely characterized. Here, we generated the first single-cell transcriptomic atlas of the *E. multilocularis* metacestode, resolving 26 transcriptionally distinct cell populations. The atlas recovered the major cell types previously described in the germinal layer, including germinative, tegumental, muscle, neuronal and putative storage cells, and revealed substantial molecular heterogeneity within several of these compartments. In particular, germinative cells segregated into distinct transcriptional states, ranging from a population enriched in markers associated with an undifferentiated germinative state to populations displaying early tegumental- or muscle-associated transcriptional programs. Notably, one of these states was strongly enriched in an isolate retaining the capacity for brood capsule and protoscolex formation but was nearly absent from a developmentally deficient isolate, suggesting a possible association between germinative-cell heterogeneity and developmental competence. Differentiated populations likewise displayed distinct molecular specializations, including developmental signaling and extracellular-matrix programs in muscle cells, microtubule-associated and transporter expression in tegumental populations, and metabolic specialization in putative storage cells. Spatial validation by whole-mount *in situ* hybridization, EdU labeling and immunofluorescence established molecular markers for major cell populations and revealed stage-specific expression patterns between metacestodes and protoscoleces. Together, these data uncover an unexpected level of molecular and cellular heterogeneity within the morphologically simple metacestode germinal layer and establish a cell-resolved framework for investigating stem-cell organization, differentiation and developmental plasticity in this medically important parasite.

## Introduction

Alveolar echinococcosis (AE) is a severe chronic disease caused by the metacestode larva of the tapeworm *Echinococcus multilocularis*, a zoonotic parasite of increasing public health concern throughout the Northern Hemisphere (Deplazes et al., 2017; Thompson, 2017). The parasite is maintained predominantly in a sylvatic transmission cycle involving foxes as definitive hosts and small rodents as intermediate hosts. Humans become accidental intermediate hosts through ingestion of parasite eggs shed in the feces of infected definitive hosts (Kern et al., 2017). Following infection, the oncosphere develops into the metacestode larva, which proliferates indefinitely through continuous asexual growth, forming an infiltrative network of vesicles that progressively invades host tissues (primarily the liver), ultimately causing AE (Kern et al., 2017; Casulli et al., 2019).

Morphologically, the metacestode is a multivesicular structure delimited by an outer acellular and carbohydrate-rich laminated layer and an inner germinal layer containing parasite cells (Thompson, 2017). The extraordinary developmental capacity of this larval stage relies on a unique population of somatic stem cells, termed germinative cells, which constitute the only proliferative cells of the larval stage and contribute to its metastatic potential. These cells generate all differentiated cell types of the metacestode, including tegumental, muscle, glycogen/lipid-storing, and nerve cells, and give rise to brood capsules and protoscoleces during development (Koziol et al., 2014; Brehm and Koziol, 2017).

Because AE is frequently diagnosed at advanced stages, complete surgical resection is often impossible, and benzimidazole chemotherapy remains the only treatment option for many patients (Brunetti et al., 2010; Hemphill et al., 2014). Although clinically effective at suppressing parasite growth, benzimidazoles are not parasiticidal, and the persistence of highly regenerative germinative cells may contribute to parasite survival and renewed growth following treatment discontinuation (Schubert et al., 2014; Brehm and Koziol, 2014; Koziol and Brehm, 2015). Deciphering the cellular and molecular mechanisms underlying the remarkable developmental capacity of the metacestode is therefore essential for developing more effective therapies. This requires a comprehensive characterization of the parasite’s cellular organization and the transcriptional programs underlying cell identity and function, particularly within the germinative cell compartment.

Over the years, considerable progress has been made toward understanding the biology of the *E. multilocularis* metacestode. Classical ultrastructural studies established the principal differentiated cell types of the larva (Sakamoto and Sugimura, 1970; Lascano et al., 1975; Koziol et al., 2013), while subsequent molecular studies identified distinctive features of *E. multilocularis* germinative cells and revealed heterogeneity within this population, suggesting the existence of distinct subpopulations with different self-renewal and differentiation capacities (Tsai et al., 2013; Koziol et al., 2014; Koziol and Brehm, 2015; Koziol et al., 2015).

More recently, bulk transcriptomic analyses identified genes associated with germinative cell function and substantially expanded the repertoire of candidate germinative-cell markers, providing new insights into the molecular programs operating within these cells (Herz et al., 2024; Herrmann et al., 2026). However, because bulk approaches average gene expression across heterogeneous cell populations, they cannot resolve the transcriptional identities of individual cell types or define the molecular organization of the metacestode at cellular resolution.

Recent advances in single-cell transcriptomics offer a powerful means to overcome these limitations by enabling unbiased molecular characterization of individual cells at high resolution (Stuart and Satija, 2019; Lähnemann et al., 2020). In flatworms, these approaches have revealed previously unrecognized cellular diversity, refined the classification of stem-cell populations, and provided molecular insights into cell differentiation, development, and tissue organization (van Wolfswinkel et al., 2014; Molinaro and Pearson, 2016; Wang et al., 2018; Fincher et al., 2018; Plass et al., 2018; Swapna et al., 2018; Zeng et al., 2018; Wendt et al., 2020; Diaz Soria et al., 2020; Li et al., 2021; Diaz Soria et al., 2024; Attenborough et al., 2024; Puckelwaldt et al., 2026). Comparable studies are, however, still lacking for cestodes, leaving the cellular composition and molecular organization of the *E. multilocularis* metacestode largely unresolved.

Here, we generated the first single-cell transcriptomic atlas of the *E. multilocularis* metacestode by combining droplet-based single-cell RNA sequencing (scRNA-seq) with extensive spatial validation using whole-mount *in situ* hybridization (ISH), EdU labelling, and immunofluorescence. Our analyses define the principal differentiated cell populations of the larval stage, establish a comprehensive repertoire of experimentally validated molecular markers, and resolve the previously recognized molecular heterogeneity of the germinative compartment into discrete transcriptional states. Together, these findings establish a molecular framework for understanding the cellular organization of the *E. multilocularis* metacestode and provide a reference atlas for future studies of parasite development, stem cell biology, and therapeutic target discovery.

## Materials and methods

### Ethics statement

Animal experiments were approved by the Ethics Committee of the Government of Lower Franconia (Regierung von Unterfranken; permit no. RUF-55.2.2-2532-2-1824-10), Würzburg, Germany, and conducted in accordance with German and European regulations on animal welfare and the institutional guidelines for the care and use of laboratory animals. All efforts were made to minimize animal suffering.

### Parasite material and *in vitro* cultivation

Parasite isolates H95 and LS23 were maintained by serial intraperitoneal passage in Meriones unguiculatus as previously described (Spiliotis and Brehm, 2009), and parasite tissue recovered from infected gerbils was used to generate *in vitro* metacestode vesicles. At the time of the experiments, H95 was unable to produce protoscoleces, whereas LS23 retained this developmental capacity. H95 metacestode vesicles were produced under standard co-culture conditions with rat Reuber hepatoma feeder cells, whereas LS23 metacestode vesicles were produced under axenic culture conditions to prevent brood capsule formation (Spiliotis and Brehm, 2009). These culture conditions were used to obtain comparable metacestode-stage material from both isolates. Vesicles from each isolate were processed independently for primary cell preparation and scRNA-seq.

### Single-cell tissue dissociation and fluorescence-activated cell sorting (FACS)

Primary cell suspensions were prepared from *in vitro*-generated metacestode vesicles using a modified version of the protocol described by Spiliotis and Brehm (2009). Briefly, metacestodes from three culture bottles were pooled, washed with phosphate-buffered saline (PBS), gently collapsed by passage through a 10 mL pipette, and incubated for 30 min at 37°C in a digestion solution containing 0.05% trypsin (Sigma T4799) and 0.02% EDTA in PBS. Following enzymatic digestion, the suspension was gently agitated and passed through a 30 μm cell strainer using a 10 mL pipette. The material retained on the strainer was recovered, resuspended in PBS, and subjected to two additional rounds of agitation and filtration. The pooled cell suspension was centrifuged at 80 × g for 1 min to remove calcareous corpuscles, after which the recovered cells were pelleted by centrifugation at 600 × g for 10 min. Cell pellets from each isolate were resuspended in sterile 1% BSA and passed through a 40 μm cell strainer (Greiner 542040). Cell suspensions were incubated with 10 U/mL RQ1 DNase (Promega M6101) for 10 min at room temperature, centrifuged at 300 × g for 5 min, and resuspended in 1% BSA. Cells were stained with 1 μL/mL eFluor 780 (Invitrogen 65-0865-14) for 20 min at room temperature in the dark to identify non-viable cells. Finally, live (eFluor 780−) cells were enriched by fluorescence-activated cell sorting using a BD FACSAria™ Fusion cell sorter and collected into microcentrifuge tubes.

### 10X Genomics library preparation and sequencing

Single-cell suspensions (∼1,500 live cells/μL) prepared from metacestode vesicles were loaded according to the standard protocol of the Chromium Single Cell 3’ Reagent Kits to capture ∼ 20000 cells per reaction. After single-cell library construction, they were sequenced on an Illumina NextSeq 2000 using one sequencing lane per sample.

### Mapping and quantification of single-cell RNA-seq

Single-cell RNA-seq data were mapped to the *Echinococcus multilocularis* reference genome EMULTI002 (GCA_000469725.3; BioProject PRJEB122) using the 10x Genomics Cell Ranger pipeline (v8.0.1). Cell-containing droplets were identified using the default Cell Ranger cell-calling algorithm. Across the two libraries, 2.07 billion sequencing reads were generated, with an average of 259,000 reads per cell. Approximately 40.6% of reads mapped confidently to the transcriptome. A total of 7,988 cells were recovered, with a median of 1,430 genes detected per cell.

### Quality control and cell clustering

The Seurat package (v5.3.0) was used to analyse the filtered counts matrices produced by Cell Ranger. Cells that had greater than 20,000 Unique Molecular Identifiers (UMIs), less than 400 genes per cell, or more than 2.5% mitochondrial transcripts were removed. Mitochondrial genes were identified from the *E. multilocularis* WormBase ParaSite WBPS19 genome annotation by selecting genes located on the mitochondrial chromosome (pathogen_EmW_mitochondrion). Following quality control, a total of 7,618 high-quality cells (3,181 H95 and 4,437 LS23 cells) were retained for downstream analyses.

Each individual dataset was normalized (NormalizeData) and variable features were identified (FindVariableFeatures, selection.method = "vst", nfeatures = 2000). Integration anchors were identified (FindIntegrationAnchors) and the two datasets were integrated (IntegrateData). The integrated expression matrix was subsequently scaled (ScaleData) and subjected to principal component analysis (PCA). The number of principal components (25) used for this analysis was defined by visual inspection of the ElbowPlot. Clusters were generated using FindNeighbors and FindClusters (resolution = 1), resulting in 26 transcriptionally distinct cell populations. The resolution for stable clustering was chosen using clustree. Low-dimensional visualization of the integrated dataset was performed using Uniform Manifold Approximation and Projection (UMAP).

### Marker identification and cluster annotation

Cluster-specific marker genes were identified using Seurat’s FindAllMarkers function with both the Wilcoxon rank-sum test (only.pos = TRUE, min.pct = 0.25, logfc.threshold = 0.25) and receiver operating characteristic (ROC) analysis (only.pos = TRUE, return.thresh = 0). Differential expression statistics obtained with the Wilcoxon test were considered together with AUC values from the ROC analysis to prioritize informative markers for cluster annotation. Final cell-type annotation was based on these analyses in combination with previously reported *E. multilocularis* markers and conserved molecular markers from other metazoans when parasite-specific markers were unavailable. Candidate genes for experimental validation were selected based on their enrichment within each cell population, expression specificity, and biological relevance.

### Gene Ontology enrichment analysis

Gene Ontology (GO) annotations for *Echinococcus multilocularis* were obtained from WormBase ParaSite. GO enrichment analysis was performed using the topGO package with the weight01 algorithm and Fisher’s exact test. Analyses were restricted to the Biological Process (BP) ontology using a minimum node size of 5. For each broad cell type, positively differentially expressed genes identified by the Wilcoxon test were used as the target gene set, whereas all expressed genes detected in the RNA assay were used as the background. P-values were adjusted for multiple testing using the Benjamini–Hochberg method, and GO terms with FDR < 0.05 were considered significantly enriched.

### Differential gene expression analysis between cell subtypes

Subtype-specific marker genes were identified using Seurat’s FindAllMarkers function (test.use = "wilcox", only.pos = TRUE, min.pct = 0.25, logfc.threshold = 0.25) by comparing each subtype with the remaining subtypes within the corresponding lineage.

### Isolate-specific stem cell subclustering

To evaluate isolate-specific heterogeneity within the germinative compartment, Stem1–Stem5 populations were extracted independently from the H95 and LS23 datasets and reanalyzed separately using Seurat. Each subset was normalized (NormalizeData), variable features were identified (FindVariableFeatures, selection.method = "vst", nfeatures = 2000), and the data were scaled prior to principal component analysis (PCA). The first 12 principal components, selected by visual inspection of the ElbowPlot, were used to construct the shared nearest-neighbor graph (FindNeighbors). Subclusters were identified using FindClusters (resolution = 0.4), with the final resolution selected based on cluster stability assessed using clustree. Low-dimensional visualization was performed using Uniform Manifold Approximation and Projection (UMAP). Subcluster-specific marker genes identified with the Wilcoxon rank-sum test (only.pos = TRUE, min.pct = 0.25, logfc.threshold = 0.25) were used to characterize the transcriptional identity of each stem cell subcluster.

### *in situ* hybridization, EdU labeling and immunofluorescence

Whole-mount fluorescent ISH was performed on *in vitro*-cultivated metacestode vesicles and activated protoscoleces according to a previously established protocol (Koziol et al., 2014). Digoxigenin (DIG)- or fluorescein-labeled antisense RNA probes were synthesized by *in vitro* transcription using T7 or SP6 RNA polymerase (New England Biolabs) and the DIG RNA Labeling Kit or Fluorescein RNA Labeling Kit (Roche) from PCR-amplified gene fragments cloned into the pJET1.2 vector (Thermo Fisher Scientific). Amplification primers for all probes used in this study are listed in Table S1. Single WMISH experiments were detected using a horseradish peroxidase (HRP)-conjugated anti-DIG antibody (Roche) followed by tyramide signal amplification (TSA) with fluorescein-tyramide. For double WMISH experiments, DIG- and fluorescein-labeled probes were hybridized simultaneously and detected sequentially using HRP-conjugated anti-fluorescein and anti-DIG antibodies (Roche), with peroxidase inactivation between detection rounds. Fluorescent signals were developed using fluorescein-tyramide and rhodamine-tyramide. EdU labeling was performed by incubating metacestode vesicles or protoscoleces with 50 μM 5-ethynyl-21-deoxyuridine (EdU; Life Technologies, Darmstadt, Germany) for 5 h or overnight, respectively. For pulse-chase experiments, EdU-labeled vesicles were returned to standard culture conditions for the indicated chase periods before fixation. EdU incorporation was detected using the Click-iT™ EdU Alexa Fluor™ 555 Imaging Kit (Thermo Fisher Scientific) according to the manufacturer’s instructions after completion of the WMISH procedure (Koziol et al., 2014; Herz et al., 2024). Where indicated, WMISH was combined with immunofluorescence using anti-Tropomyosin or anti-FMRFamide antibodies as previously described (Koziol et al., 2013; Koziol et al., 2016a). Nuclei were counterstained with DAPI. Fluorescence images were acquired using a Nikon Eclipse Ti2-E confocal microscope and processed with Fiji/ImageJ as previously described (Herz et al., 2024). In all experiments, labeled sense probes were used as negative controls and yielded no detectable signal.

## Results

### Single-cell transcriptomic profiling defines the cellular landscape of the *Echinococcus multilocularis* metacestode

To comprehensively characterize the cellular composition of the *E. multilocularis* metacestode, we performed scRNA-seq on two independent samples of *in vitro*-generated metacestode vesicles (Fig. 1A). Parasite tissue was dissociated using trypsin and viable cells were enriched by fluorescence-activated cell sorting prior to library preparation using the 10x Genomics Chromium platform. A total of 7,988 cells were sequenced across both samples, of which 7,618 passed quality-control filtering and were retained for downstream analyses (Table S2).

**Fig. 1.**
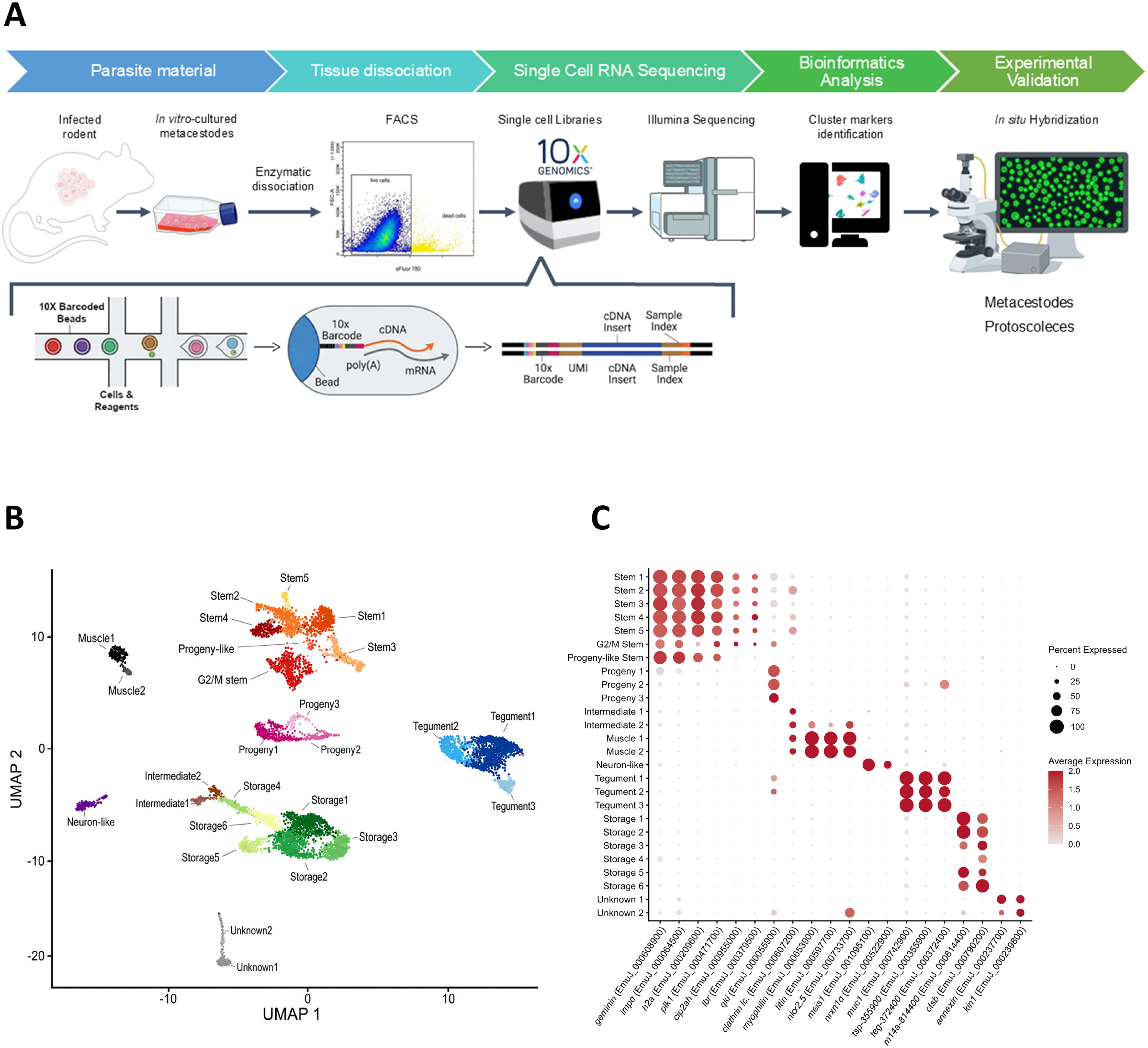
Single-cell transcriptomic atlas reveals the major cell populations of the *Echinococcus multilocularis* metacestode. (A) Overview of the experimental workflow. *in vitro*-generated metacestode vesicles were enzymatically dissociated, and viable cells (eFluor780-negative) were enriched by fluorescence-activated cell sorting (FACS) prior to single-cell RNA sequencing using the 10x Genomics Chromium 3□ platform. Unsupervised clustering was used to identify transcriptionally distinct cell populations, and population-specific marker genes were selected as candidates for experimental validation by whole-mount *in situ* hybridization. (B) Uniform Manifold Approximation and Projection (UMAP) of 7,618 high-quality metacestode cells showing the 26 transcriptionally distinct cell populations identified in the integrated atlas. (C) Dot plot showing the expression of selected population-specific marker genes representing the major cell populations. These markers were selected as candidates for experimental validation of cell identity. Dot size indicates the percentage of cells expressing each marker within a cluster, whereas color intensity represents the average normalized expression level.

Dimensionality reduction followed by unsupervised clustering resolved 26 transcriptionally distinct cell populations (Fig. 1B, Table S3). Cell identities were inferred using previously characterized *E. multilocularis* markers where available (Koziol et al., 2014; Herz et al., 2024) and conserved molecular markers from other metazoans when parasite-specific markers were lacking.

The resulting cellular atlas comprised six stem cell clusters, three tegumental clusters, six storage cell clusters, two muscle cell clusters, and one neuron-like cluster. In addition, we identified three small progeny clusters with transcriptional profiles distinct from both germinative cells and fully differentiated populations. Two additional intermediate clusters exhibited partial expression of genes associated with differentiated lineages while lacking the complete molecular signatures of mature cell types. Finally, two clusters could not be confidently assigned to any previously described cell type (Fig. 1B). We also identified a small progeny-like stem cluster that retained a stem cell-associated transcriptional program despite reduced expression of several canonical germinative cell markers.

To further validate these annotations, we performed Gene Ontology (GO) enrichment analysis using Biological Process terms (Fig. S1). The resulting functional signatures were highly consistent with the inferred identities of each major cell population. Stem cell clusters showed strong enrichment for processes associated with cell proliferation and genome maintenance, reflecting their highly proliferative nature. Storage cells were predominantly enriched for metabolic pathways, consistent with their inferred metabolic functions. Tegumental cells displayed significant enrichment for microtubule-based processes, in agreement with the specialized cytoskeletal organization required for tegument formation and maintenance. Interestingly, muscle cells were enriched for Wnt signaling, a pathway widely implicated in developmental patterning across metazoans, consistent with previous studies proposing developmental and morphogenetic functions for these cells in the parasite (Koziol et al., 2016a). Overall, the concordance between transcriptional identity and functional enrichment provides independent support for the inferred identities of the major cell populations.

To experimentally validate these annotations, we selected candidate marker genes based on their transcriptomic enrichment and expression patterns within each major cell population (Fig. 1C, Table S3, Table S4) and examined their spatial expression by whole-mount ISH in both metacestode vesicles and protoscoleces.

### Novel markers of the germinative cell population

To identify molecular markers of the germinative cell population defined by scRNA-seq, we searched for transcripts consistently enriched across stem cell clusters. In addition to the established stem cell marker *cip2ah* (EmuJ_000955000) (Herz et al., 2024), we identified multiple genes whose expression was largely restricted to germinative cells (Fig. 1C, Table S3). We selected histone 2A (h2a, EmuJ_000209600), importin subunit alpha (impα, EmuJ_000064500), and lamin B receptor (*lbr*; EmuJ_000379500) for experimental validation.

As an initial validation, we examined the expression of these candidate markers in combination with EdU labeling, which specifically labels cells undergoing DNA replication. As expected for markers of the germinative cell population, virtually all EdU-positive cells expressed *lbr*, h2a, or impα following a 5 h EdU pulse, whereas only a subset of marker-positive cells incorporated EdU (Fig. 2A, Fig. S2A). *lbr* showed a similar association with EdU-positive cells in activated protoscoleces (Fig. 2B), indicating that its expression is conserved across larval forms.

**Fig. 2.**
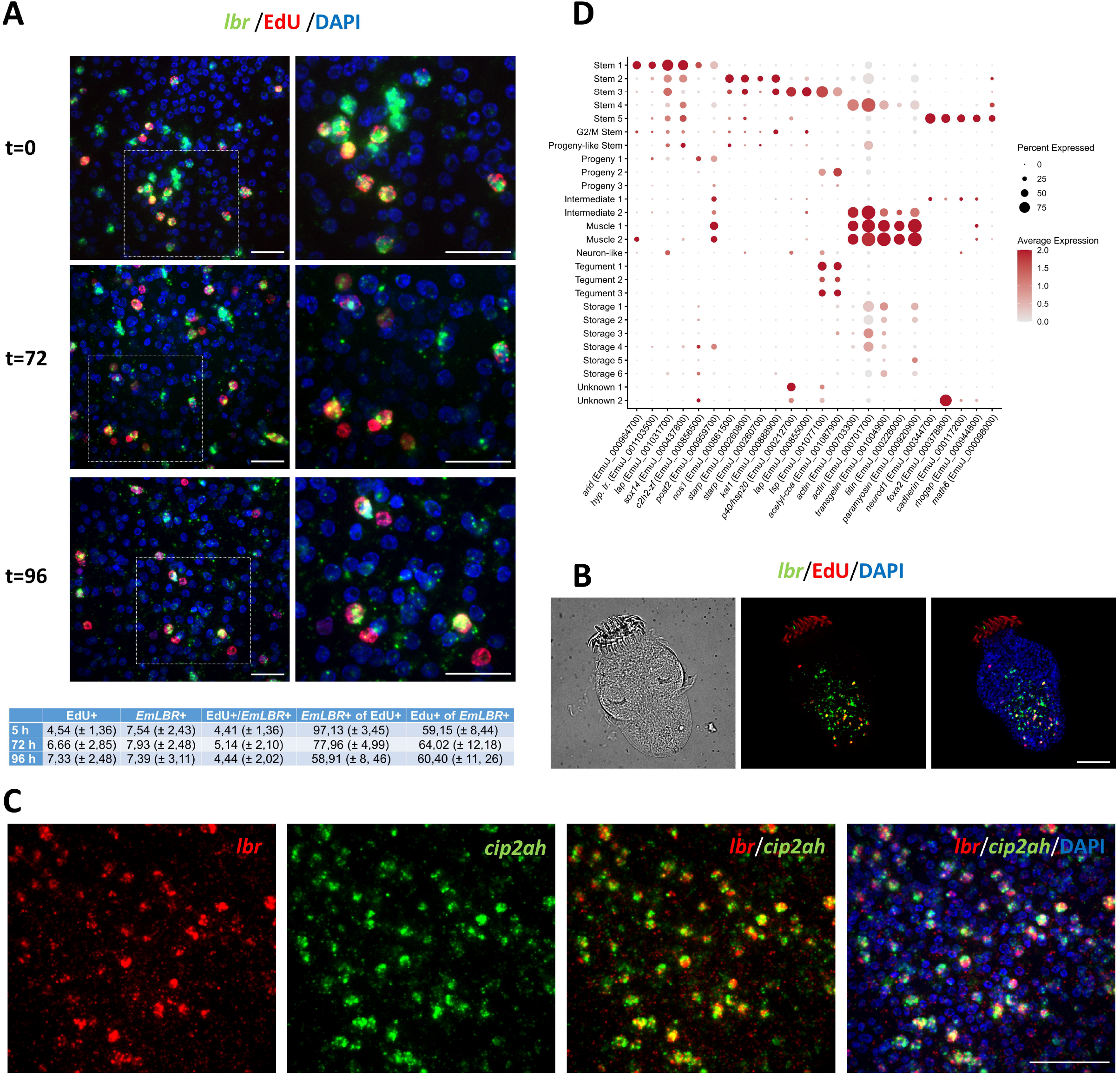
Experimental validation of *lbr* as a germinative cell marker and identification of molecular signatures defining stem cell populations. (A) Validation of *lbr* expression by whole-mount *in situ* hybridization (green) combined with EdU labeling (red) in metacestode vesicles. Parasites were exposed to EdU for 5 h and analyzed immediately after the pulse (t = 0) or following 72 h and 96 h of chase. Nuclei were counterstained with DAPI (blue). Dashed boxes are magnified in the images to the right. Quantification of EdU labeling, *lbr* expression, double-positive cells, and the proportions of EdU-positive cells expressing *lbr* and *lbr*-positive cells incorporating EdU is shown below the corresponding images. Scale bar, 20 µm. (B) Combined detection of *lbr* expression by whole-mount *in situ* hybridization (green) and EdU incorporation (red) in activated protoscoleces. Nuclei were counterstained with DAPI (blue). Scale bar, 40 µm. (C) Double fluorescence whole-mount *in situ* hybridization showing co-expression of *lbr* (red) and the established germinative cell marker *cip2ah* (green) in metacestode vesicles. Merged images are shown with and without DAPI nuclear staining (blue). Scale bar, 40 µm. (D) Dot plot showing the expression of selected marker genes distinguishing the stem cell populations identified in the integrated atlas. Marker genes were selected based on their population-specific expression patterns. Dot size indicates the percentage of cells expressing each marker within each population, whereas color intensity indicates the average normalized expression level.

We next asked whether *lbr* expression is maintained following commitment to differentiation. To address this, we performed EdU pulse-chase experiments and analyzed metacestodes 72 or 96 h after a 5 h EdU pulse. Because differentiating progeny retain the EdU label while progressively losing germinative cell identity, the proportion of *lbr*+/EdU+ cells was expected to decline if *lbr* expression were restricted to germinative cells. Consistent with this prediction, the fraction of *lbr*+/EdU+ cells decreased progressively throughout the chase period (Fig. 2A).

As an independent experimental validation, we performed double whole-mount ISH using the established germinative cell marker *cip2ah*. *lbr*, h2a, and impα each exhibited extensive co-localization with *cip2ah* within the germinal layer (Fig. 2C, Fig. S2B, Fig. S2C), confirming that they label the same germinative cell population.

Together, these complementary approaches establish *lbr*, h2a, and impα as robust molecular markers of the *E. multilocularis* germinative cell population.

### Transcriptomic heterogeneity of the germinative compartment

Beyond the proliferative transcriptional program shared by all germinative cells, unsupervised clustering resolved six molecularly distinct stem cell populations. One population displayed a transcriptional profile consistent with actively proliferating cells in the G2/M phase of the cell cycle and was therefore designated G2/M Stem. The remaining populations exhibited specific molecular signatures consistent with different transcriptional states within the germinative compartment (Fig. 2D, Table S5).

Among these, Stem2 displayed the strongest transcriptional signature of an undifferentiated germinative state. Its most prominent markers included nos1 (EmuJ_000861500) and two STARP-like antigens (EmuJ_000260800 and EmuJ_000260700). nos1, the *E. multilocularis* orthologue of the conserved stem cell regulator nanos, has previously been reported to be expressed in a small subpopulation of germinative cells within the germinal layer (Koziol et al., 2014). Stem2 was also enriched for kal1 (EmuJ_000888900), recently identified as a marker of a slowly cycling germinative cell population (Herz et al., 2024). The enrichment of these established germinative cell markers, together with the absence of lineage-associated transcriptional programs, is consistent with Stem2 representing the least lineage-committed population identified in the atlas.

In contrast, Stem1 was distinguished by a regulatory program characterized by the coordinated expression of the chromatin-associated regulator arid (EmuJ_000964700), the transcription factors SOX14 (EmuJ_000437800) and the posterior Hox gene Post2 (EmuJ_000959700), together with a C2H2-type zinc finger protein (EmuJ_000856500), a hypothetical transcript (EmuJ_001103500), and a leucyl aminopeptidase family member (EmuJ_001031700). Together, these genes define a distinct transcriptional program in Stem1. Unlike Stem2, however, Stem1 lacked the canonical germinative markers nos1 and kal1 and did not exhibit transcriptional signatures associated with any differentiated lineage identified in the atlas. These observations are consistent with Stem1 representing a molecularly distinct stem cell state whose lineage relationships remain unresolved.

Similarly, Stem5 was enriched for developmental regulators, including neurod1 (EmuJ_000344700), foxa2 (EmuJ_000378800), math6 (EmuJ_000098000), together with cadherin (EmuJ_000117200) and a rhogap (EmuJ_000944800). Although neurod1 is broadly associated with neuronal differentiation across metazoans (Scimone et al., 2014; Wendt et al., 2020), these genes were strongly enriched in Stem5 and showed little expression in differentiated neuron-like cells (Fig. 2D), indicating that this population represents a distinct regulatory state rather than a neuronal lineage.

The remaining stem cell populations displayed transcriptional programs consistent with lineage priming. Stem3 was characterized by high expression of a tetraspanin (EmuJ_001077100), together with an acetyl-CoA hydrolase (EmuJ_001087900), a putative p40/hsp20 protein (EmuJ_000212700), and a leucyl aminopeptidase family member (EmuJ_000855000). Most of these genes were largely restricted to Stem3, whereas the tetraspanin and acetyl-CoA hydrolase were also expressed in differentiated tegumental cells (Fig. 2D). In contrast, canonical tegumental markers, including muc-1 (EmuJ_000742900), were largely absent from Stem3 and restricted to differentiated tegumental populations (Fig. 1C).

This transcriptional profile is therefore consistent with a stem cell population exhibiting early tegumental lineage priming. Similarly, Stem4 displayed a transcriptional program enriched in genes associated with cytoskeletal organization and contractility, including two actin isoforms (EmuJ_000703300 and EmuJ_000701700), transgelin (EmuJ_001004900), titin (EmuJ_000226000), and paramyosin (EmuJ_000920900). These genes were also highly expressed in differentiated muscle cells, indicating that Stem4 shares much of its transcriptional program with the muscle lineage. This expression profile is consistent with a muscle-lineage-primed progenitor population.

Collectively, these analyses reveal substantial molecular heterogeneity within the germinative compartment. Stem2 exhibits the strongest transcriptional signature of an undifferentiated germinative state, whereas Stem3 and Stem4 display gene expression programs consistent with early tegumental and muscle lineage priming. Stem1 and Stem5 are defined by distinct regulatory programs that cannot yet be assigned to specific differentiation trajectories. Together, these findings reveal multiple transcriptionally distinct stem cell populations and provide a framework for future studies aimed at resolving their developmental relationships.

### Isolate-specific differences in the germinative compartment

Having established the molecular heterogeneity of the germinative compartment, we examined whether the identified stem cell states were similarly represented in the two biological samples used to generate the metacestode cell atlas. Although both samples were integrated into a common atlas, they originated from parasite isolates with different developmental competence. The H95 isolate has irreversibly lost the ability to generate protoscoleces in vivo, whereas the LS23 isolate retains this capacity but was maintained under axenic culture conditions to prevent brood capsule formation. Consequently, both samples represented comparable metacestode-stage material, providing an opportunity to assess whether developmental competence is associated with differences in stem cell composition.

Examination of the atlas revealed one notable difference between the two samples. Stem3 was well represented in LS23 but was detected as only a single cell in H95, constituting the most pronounced difference in stem cell composition between the two isolates.

To investigate this observation further, stem cell populations (Stem1–Stem5) were extracted from each isolate and reanalyzed independently. Subclustering of H95 stem cells resolved four populations corresponding to Stem1-, Stem2-, Stem4-, and Stem5-like states, whereas no discrete Stem3-like cluster could be resolved (Table S6). In contrast, six stem cell populations were identified in LS23 (Table S7). Comparison with the transcriptional states defined in the integrated atlas showed that four corresponded to Stem1-, Stem2-, Stem4-, and Stem5-like populations, whereas the remaining two corresponded to two transcriptionally distinct Stem3-like states. Both LS23 Stem3-like populations retained the core transcriptional features that define the Stem3 population identified in the integrated atlas. One closely recapitulated the Stem3 transcriptional program, characterized by strong expression of the Stem3-specific tetraspanin marker (EmuJ_001077100) together with acetyl-CoA hydrolase (EmuJ_001087900), leucyl aminopeptidase (EmuJ_000855000), and p40/hsp20 (EmuJ_000212700). The second retained elements of this Stem3 signature while additionally exhibiting prominent expression of noggin (EmuJ_000089300), npp27 (EmuJ_000347700), neuroendocrine convertase 2 (EmuJ_000412100), GATA2 binding factor 2 (EmuJ_000215900), and other regulatory genes (Table S7), revealing further transcriptional diversification within the Stem3 compartment rather than the emergence of a distinct stem cell class.

The exclusive detection of both Stem3-like populations in LS23 raises the possibility that they are associated with developmental competence for brood capsule and protoscolex formation. However, the developmental relationship between these populations remains to be established experimentally.

Because fewer stem cells were recovered from H95 than from LS23, the apparent underrepresentation of Stem3-derived populations in H95 should be interpreted with caution. Although these observations are consistent with biological differences between the two isolates, reduced sampling depth may also have limited the detection of rare stem cell states.

### Identification and validation of muscle cell markers

The two muscle populations identified in the transcriptomic atlas were defined by the expression of established muscle-associated genes, including paramyosin (EmuJ_000920900) and a myosin essential light chain (EmuJ_000835500). Among the most robust molecular markers shared by both populations were titin (EmuJ_000597700) and myophilin (EmuJ_000653900), which displayed strong and highly specific expression across nearly all muscle cells while remaining largely absent from other cell types (Fig. 1C).

Whole-mount ISH combined with EdU labeling confirmed that neither titin nor myophilin was expressed in the proliferative germinative cell population (Fig. 3A). In both cases, labeled cells displayed a characteristic morphology of muscle cells, with cell bodies giving rise to long, branched cytoplasmic processes.

**Fig. 3.**
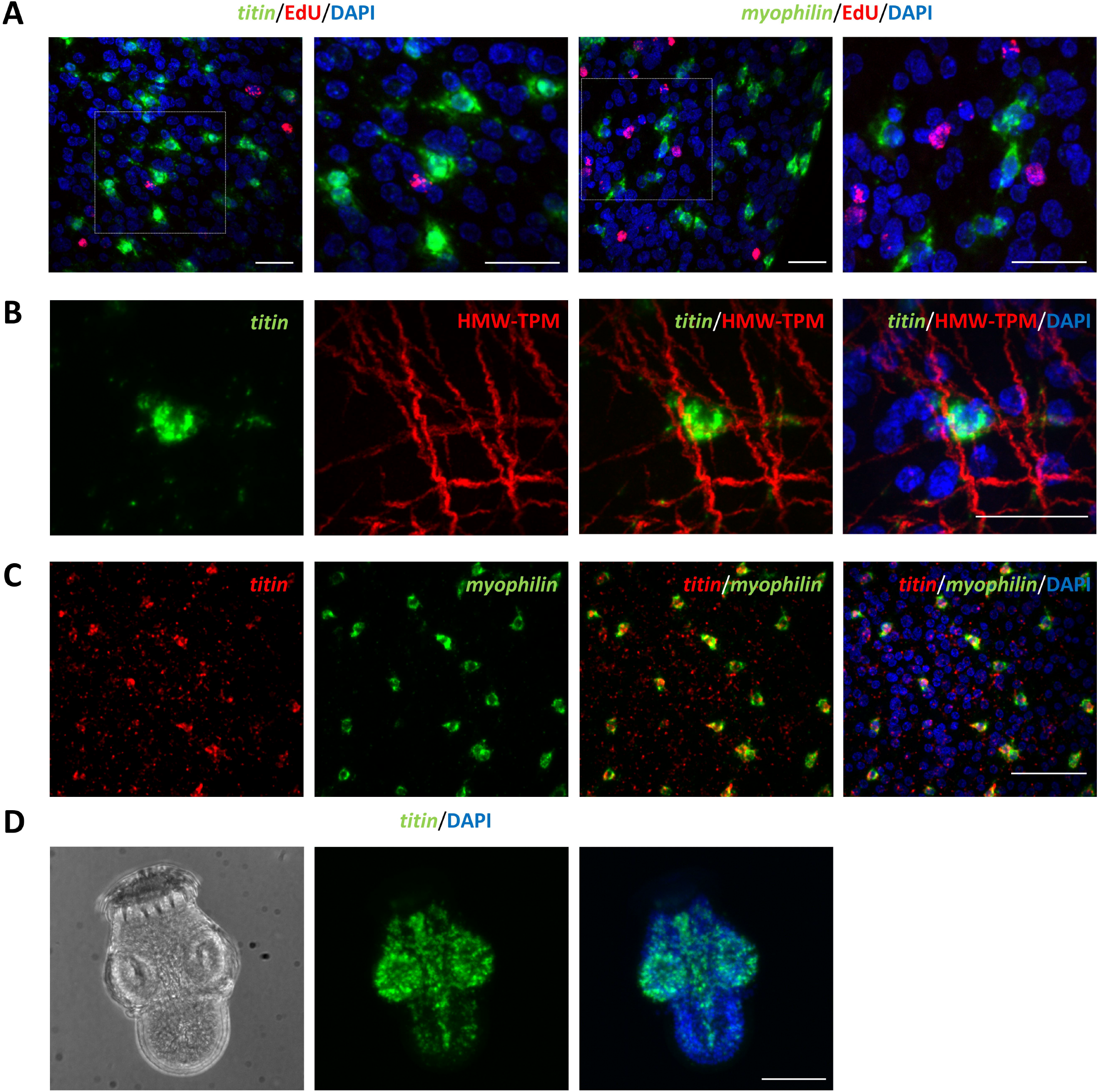
Experimental validation of *titin* and *myophilin* as molecular markers of muscle cells. (A) Whole-mount *in situ* hybridization showing *titin* (green, left) and *myophilin* (green, right) expression in metacestode vesicles combined with EdU labeling (red). Parasites were exposed to EdU for 5 h prior to fixation. Nuclei were counterstained with DAPI (blue). Dashed boxes are magnified in the images to the right. Scale bars, 20 µm. (B) Combined detection of *titin* transcripts by whole-mount *in situ* hybridization (green) and Tropomyosin immunofluorescence (red) in metacestode vesicles. Merged images are shown with and without DAPI nuclear staining (blue). Scale bar, 20 µm. (C) Double fluorescence whole-mount *in situ* hybridization showing co-expression of *titin* (red) and *myophilin* (green) in metacestode vesicles. Merged images are shown with and without DAPI nuclear staining (blue). Scale bar, 40 µm. (D) Whole-mount *in situ* hybridization showing *titin* expression in an activated protoscolex. Bright-field, fluorescence, and merged images with DAPI nuclear staining are shown. Scale bar, 40 µm.

Muscle identity was further validated by combining titin ISH with anti-Tropomyosin immunofluorescence. All observed titin-positive cells were associated with Tropomyosin-positive fibers, confirming their identity as differentiated muscle cells (Fig. 3B). Furthermore, double whole-mount ISH revealed complete co-localization between titin and myophilin, demonstrating that both markers identify the same muscle cell population (Fig. 3C).

Both markers were broadly expressed throughout the musculature of the two larval stages. In metacestodes, positive cells were distributed throughout the germinal layer (Fig. 3A, Fig. 3C), consistent with the organization of the subtegumental musculature. In protoscoleces, titin and myophilin labeled muscle cells throughout the body, including the subtegumental musculature, the suckers, the musculature surrounding the rostellum, and rows of longitudinal muscle bundles associated with scolex invagination (Fig. 3D). These expression patterns closely matched the muscular system previously described by phalloidin labeling (Koziol et al., 2013), indicating that both genes act as general markers of the larval musculature.

Together, these complementary approaches establish titin and myophilin as robust molecular markers of differentiated muscle cells in the *E. multilocularis* metacestode. Their conserved expression throughout the musculature of both larval stages further demonstrates that they constitute general molecular markers of the larval muscular system.

In addition to these broadly expressed muscle markers, we examined *nkx2.5* (EmuJ_000733700), another gene expressed in both muscle populations identified in the transcriptomic atlas (Fig. 1C). *nkx2.5* was likewise absent from EdU-positive cells and associated with Tropomyosin-positive muscle fibers in metacestodes (Fig. S3A, Fig. S3C). Double whole-mount ISH further revealed overlapping expression with titin, although the relative signal intensity of the two transcripts varied among individual cells (Fig. S3D). In activated protoscoleces, however, *nkx2.5* expression was restricted to a subset of posterior muscle cells, in contrast to the broad distribution observed for titin and myophilin (Fig. S3B). Notably, this posteriorly restricted expression pattern resembled that previously described for wnt11a (EmuJ_000907500) and wnt1 (EmuJ_000349900), two posterior Wnt ligands expressed by muscle cells in *E. multilocularis* (Koziol et al., 2016a). Consistent with these previous observations, our single-cell data confirmed that expression of these posterior Wnt ligands in the metacestode is restricted to the muscle populations. Conversely, genes associated with anterior identity, including the Wnt inhibitors sfrp (EmuJ_000838700), sfl (EmuJ_001023000) and frizzled-10 (EmuJ_000085700), as well as the FGF signaling components nou-darake (EmuJ_000770900) and nou-darake-like (EmuJ_000816800), were detected only at very low levels in the metacestode atlas. Together, these findings are consistent with the proposed broadly posteriorized identity of the metacestode, in which anterior development is suppressed until the onset of brood capsule and protoscolex formation (Koziol et al., 2016a; Herrmann et al., 2026).

Both muscle populations also exhibited a prominent extracellular matrix (ECM) transcriptional signature, characterized by the enrichment of multiple collagens (EmuJ_000140000, EmuJ_000140100, EmuJ_000823800, and EmuJ_000417600) and basement membrane-associated components, including basement membrane specific heparan sulfate (EmuJ_000701800), hemicentin (EmuJ_000422300), and fibrillins (EmuJ_000997200 and EmuJ_001054000) (Table S3).

Comparative differential expression analysis identified distinct molecular signatures for the two muscle populations (Table S8). Muscle population 1 was characterized by preferential expression of a Wnt family member (wnt11b, EmuJ_000104000), sarcomeric α-actinin (EmuJ_001096300), and anti-dorsalizing morphogenetic protein 1a (ADMP1a) (EmuJ_000181200), whereas muscle population 2 preferentially expressed an amiloride-sensitive sodium channel (EmuJ_000424600), delta-like protein (EmuJ_000744300), noggin (EmuJ_000089300), and an orphan G protein-coupled receptor (EmuJ_000672800). These genes provide candidate molecular markers for future studies aimed at determining whether these transcriptomic populations correspond to spatially or functionally distinct muscle subpopulations.

### Identification and validation of neuronal cell markers

The transcriptomic atlas identified a neuron-like cluster characterized by the expression of multiple genes involved in synaptic organization and neuronal signaling, including neurexin 1 alpha (nrx1α, EmuJ_000522900), synaptic vesicle 2 (EmuJ_000441900), ELKS/CAST (EmuJ_000962800), and a neuronal Ca²⁺ sensor (EmuJ_001111000), together with the homeobox transcription factor meis1 (EmuJ_001095100) and an intermediate filament protein (EmuJ_000858200) (Fig. 1C, Table S3). From these candidates, we selected the transcription factor meis1 and nrx1α for experimental validation.

Whole-mount ISH combined with EdU labeling showed that neither meis1 nor nrx1α was detected in EdU-positive cells. Notably, nrx1α-positive cells displayed long cellular projections consistent with neurites (Fig. 4A). In metacestode vesicles containing brood capsules, both genes were expressed throughout the germinal layer but were not detected within the brood-capsule tissue itself.

**Fig. 4.**
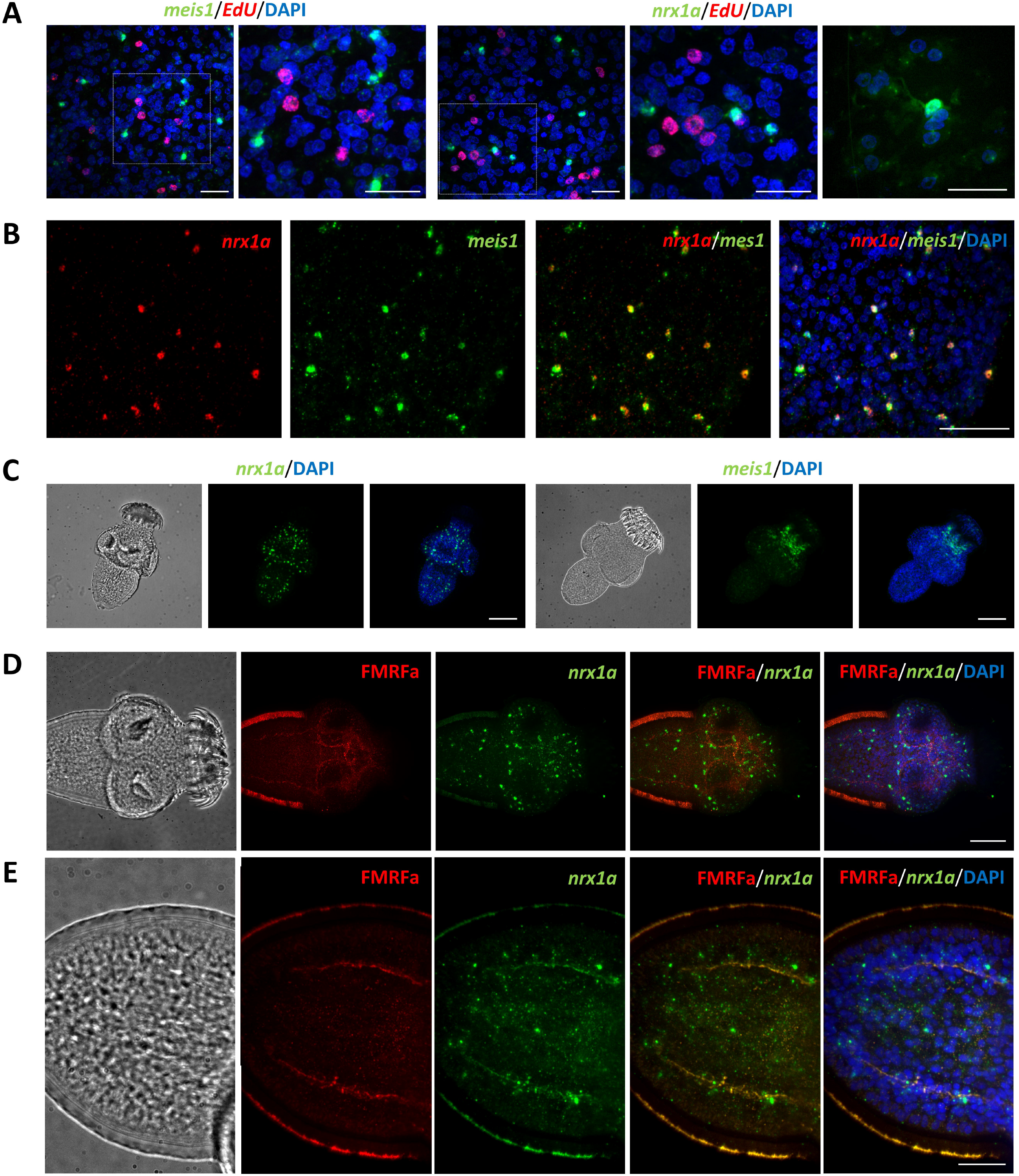
Experimental validation of *meis1* and *nrx1α* as molecular markers of neuronal cells. (A) Whole-mount *in situ* hybridization showing *meis1* (green, left) and *nrx1α* (green, right) expression in metacestode vesicles combined with EdU labeling (red). Parasites were exposed to EdU for 5 h prior to fixation. Nuclei were counterstained with DAPI (blue). Dashed boxes are magnified in the images to the right. Scale bars, 20 µm. (B) Double fluorescence whole-mount *in situ* hybridization showing co-expression of *nrx1α* (red) and *meis1* (green) in metacestode vesicles. Merged images are shown with and without DAPI nuclear staining (blue). Scale bar, 40 µm. (C) Whole-mount *in situ* hybridization showing *nrx1α* (left) and *meis1* (right) expression in activated protoscoleces. Bright-field, fluorescence, and merged images with DAPI nuclear staining are shown. Scale bar, 40 µm. (D) Combined detection of *nrx1α* transcripts by whole-mount *in situ* hybridization (green) and FMRFamide immunofluorescence (red) in activated protoscoleces. Merged images are shown with and without DAPI nuclear staining (blue). Scale bar, 40 µm. (E) Higher-magnification view of the posterior region of an activated protoscolex showing *nrx1α* expression (green) in association with FMRFamide-positive nerve cords (red). Merged images are shown with and without DAPI nuclear staining (blue). Scale bar, 20 µm.

Double whole-mount ISH revealed largely overlapping expression of *meis1* and *nrx1α* in metacestodes (Fig. 4B). In contrast, their expression patterns diverged markedly in protoscoleces. Whereas *meis1* expression was restricted to a ring surrounding the rostellum, *nrx1α* labeled a more elaborate network extending along the anterior-posterior axis of the scolex (Fig. 4C, Video S1), closely matching the organization of the protoscolex central nervous system described previously (Koziol et al., 2013; Koziol et al., 2016b).

To further validate the neuronal identity of *nrx1α*-expressing cells, *nrx1α* ISH was combined with anti-FMRFamide immunofluorescence, which provides the most comprehensive visualization of the protoscolex nervous system (Koziol et al., 2013). Although direct cellular co-localization was difficult to assess throughout the nervous system, the two signals showed closely corresponding anatomical distributions (Fig. 4D). This is consistent with the distinct labeling properties of the two approaches: *nrx1α* ISH predominantly labeled neuronal cell bodies, whereas FMRFamide immunoreactivity primarily delineated nerve fibers and tracts, in which individual neuronal cell bodies are difficult to resolve (Koziol et al., 2013). At higher magnification, however, occasional FMRFamide-positive structures resembling neuronal cell bodies could be distinguished along the main lateral nerve cords, in close spatial association with *nrx1α*-positive cells (Fig. 4E).

Together, these experiments support *meis1* and *nrx1α* as transcriptional markers of neuronal populations in *E. multilocularis*. Moreover, the distinct expression patterns of *meis1* and *nrx1α* in metacestodes and protoscoleces reveal differences in the molecular organization of neuronal populations between the two larval stages. These findings provide independent molecular support for previous developmental studies showing that the protoscolex nervous system develops independently from that of the metacestode (Koziol et al., 2013).

### Novel markers of the tegumental cell population

The tegumental cell populations identified in the transcriptomic atlas were assigned based on the expression of the previously described tegumental marker *muc1* (EmuJ_000742900) (Koziol et al., 2014). To identify additional molecular markers of differentiated tegumental cells, we searched for transcripts specifically expressed in the tegumental compartment. Among the most robust candidates were a tetraspanin (tsp-355900, EmuJ_000355900) and a tegumental protein containing a dynein light-chain domain (teg-372400, EmuJ_000372400), both of which displayed strong and highly specific expression in tegumental cells with little or no expression in other cell populations (Fig. 1C, Table S3).

Whole-mount ISH combined with EdU labeling showed that neither tsp-355900 nor teg-372400 was detected in EdU-positive cells (Fig. 5A). In metacestodes, double whole-mount ISH demonstrated extensive co-localization between the two transcripts, indicating that they label the same cell population (Fig. 5B). Whereas tsp-355900 was expressed in the metacestode germinal layer and brood-capsule wall, its expression was not detected in protoscoleces (Fig. 5E, Fig. S4A, Fig. S4B). In contrast, teg-372400 expression was already evident in developing brood capsules and was maintained in protoscoleces, with additional expression detected in the protonephridial system (Fig. 5E, Fig. S4C, Fig. S4D). These distinct expression patterns indicate stage-specific changes in tegumental gene expression during larval development.

**Fig. 5.**
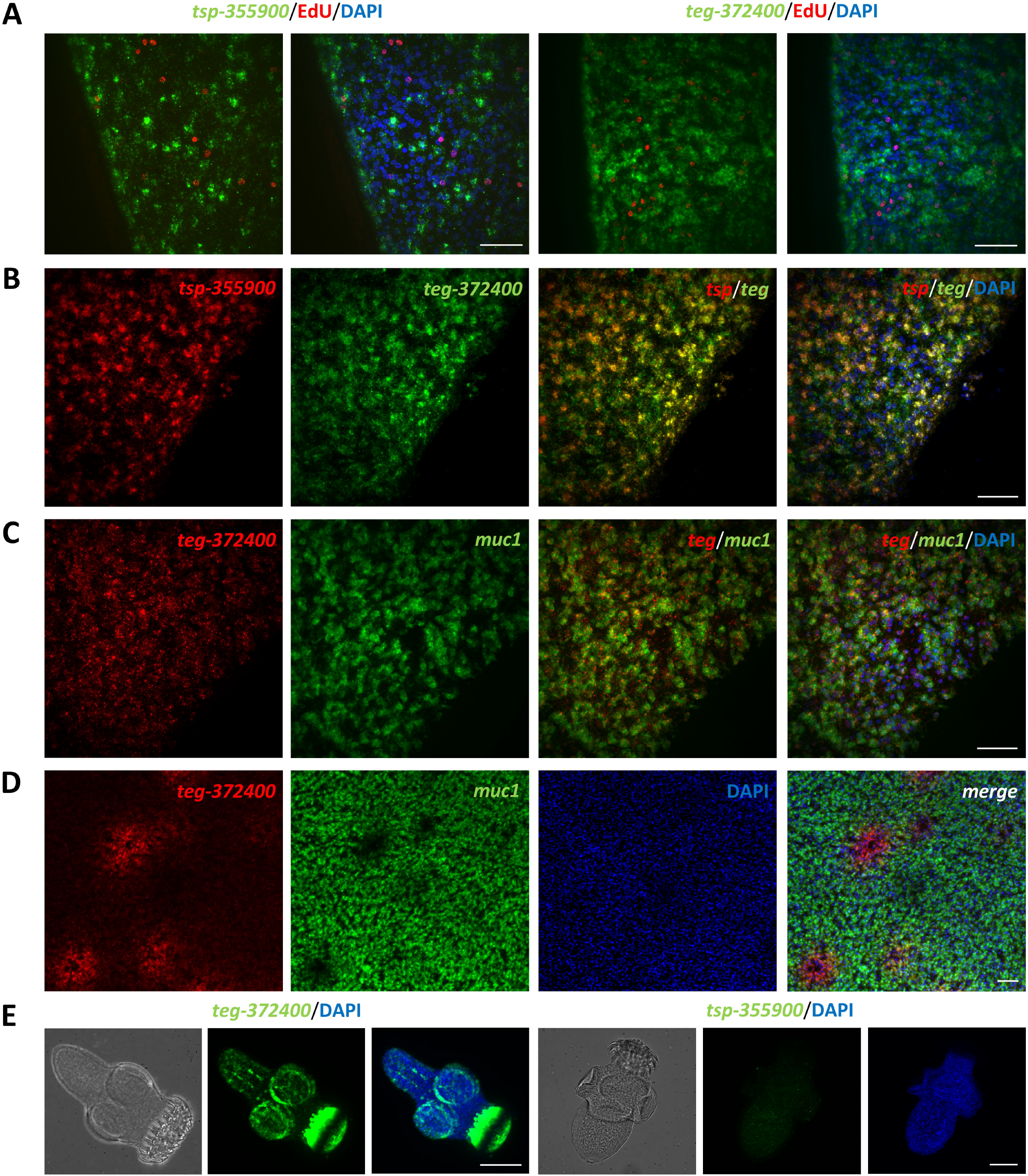
Experimental validation of *tsp-355900* and *teg-372400* as molecular markers of tegumental cells. (A) Whole-mount *in situ* hybridization showing *tsp-355900* (green, left) and *teg-372400* (green, right) expression in metacestode vesicles combined with EdU labeling (red). Parasites were exposed to EdU for 5 h prior to fixation. Nuclei were counterstained with DAPI (blue). Scale bars, 20 µm. (B) Double fluorescence whole-mount *in situ* hybridization showing co-expression of *tsp-355900* (red) and *teg-372400* (green) in metacestode vesicles. Merged images are shown with and without DAPI nuclear staining (blue). Scale bar, 40 µm. (C) Double fluorescence whole-mount *in situ* hybridization showing co-expression of *teg-372400* (red) and the previously described tegumental marker *muc1* (green) in metacestode vesicles. Merged images are shown with and without DAPI nuclear staining (blue). Scale bar, 40 µm. (D) Representative field showing localized domains with comparatively strong *teg-372400* expression and reduced *muc1* signal. Merged image includes DAPI nuclear staining (blue). Scale bar, 40 µm. (E) Whole-mount *in situ* hybridization showing *teg-372400* (left) and *tsp-355900* (right) expression in activated protoscoleces. Bright-field, fluorescence, and merged images with DAPI nuclear staining are shown. Scale bar, 40 µm.

To further validate the newly identified markers, *teg-372400* ISH was combined with *muc1* (Fig. 5C). Extensive co-localization throughout the germinal layer confirmed that the cell population labeled by *teg-372400* and *tsp-355900* corresponds to tegumental cells, thereby validating both genes as novel molecular markers of the tegument. However, discrete domains exhibiting high *teg-372400* expression together with reduced *muc1* signal were also detected (Fig. 5D). These domains lacked the characteristic nuclear organization of morphologically identifiable developing brood capsules. Given the differential expression of these markers during brood-capsule and protoscolex development, these domains may represent regions undergoing early tegumental remodeling, although their developmental fate remains unknown.

Together, these experiments establish *tsp-355900* and *teg-372400* as novel molecular markers of tegumental cells in the *E. multilocularis* metacestode. Their distinct expression patterns during brood-capsule and protoscolex development further reveal stage-specific changes in the molecular composition of the tegument.

Beyond the identification of general tegumental markers, the transcriptomic atlas also resolved three transcriptionally distinct tegumental populations. Although all three populations shared a common tegumental transcriptional program, comparative differential expression analysis identified distinct molecular signatures among them (Table S9). Tegument 1 was characterized by preferential expression of several genes encoding membrane transporters, including a solute carrier family 5 member (EmuJ_000714000), an amino acid transporter (EmuJ_001182500), and a solute carrier family 43 member (EmuJ_000763900). In contrast, Tegument 3 showed preferential expression of several dynein light-chain genes, including EmuJ_000946900 and EmuJ_000941000. Although no single gene uniquely defined each population, these distinct patterns of preferential gene expression reveal previously unrecognized molecular heterogeneity within the tegumental compartment and provide candidate markers for future studies aimed at determining whether these populations represent spatially, functionally, or developmentally distinct tegumental states.

### Identification of molecular markers associated with putative storage cells

The transcriptomic atlas identified six transcriptionally distinct populations that most likely represent storage cells based on a shared transcriptional signature enriched in genes associated with nutrient metabolism and storage, including ferritin (EmuJ_000382200), Antigen B family members (e.g., EmuJ_000381200, EmuJ_000381500), glutamine synthetase (EmuJ_000959900), fatty acid-binding proteins (EmuJ_000550000, EmuJ_000549800), and multiple enzymes involved in carbohydrate metabolism (Table S3). To identify molecular markers associated with this putative storage cell compartment, we searched for transcripts broadly expressed across these populations. Among the candidate genes, we selected cathepsin B (ctsb, EmuJ_000790200) and a previously uncharacterized M14A family peptidase (m14a-814400, EmuJ_000814400) for experimental validation.

Whole-mount ISH combined with EdU labeling showed that neither ctsb nor m14a-814400 was detected in EdU-positive cells (Fig. 6A). In metacestode vesicles, both genes labeled cells distributed throughout the germinal layer, often arranged in interconnected groups. Double whole-mount ISH revealed extensive co-localization between ctsb and m14a-814400, indicating that the two genes are expressed in largely overlapping differentiated cell populations (Fig. 6B). Both transcripts were also detected in protoscoleces, indicating that expression of these markers is maintained across larval stages. Interestingly, whereas the m14a-814400 produced a sharply defined cellular staining pattern, ctsb exhibited a more diffuse distribution within the tissue (Fig. 6C).

**Fig. 6.**
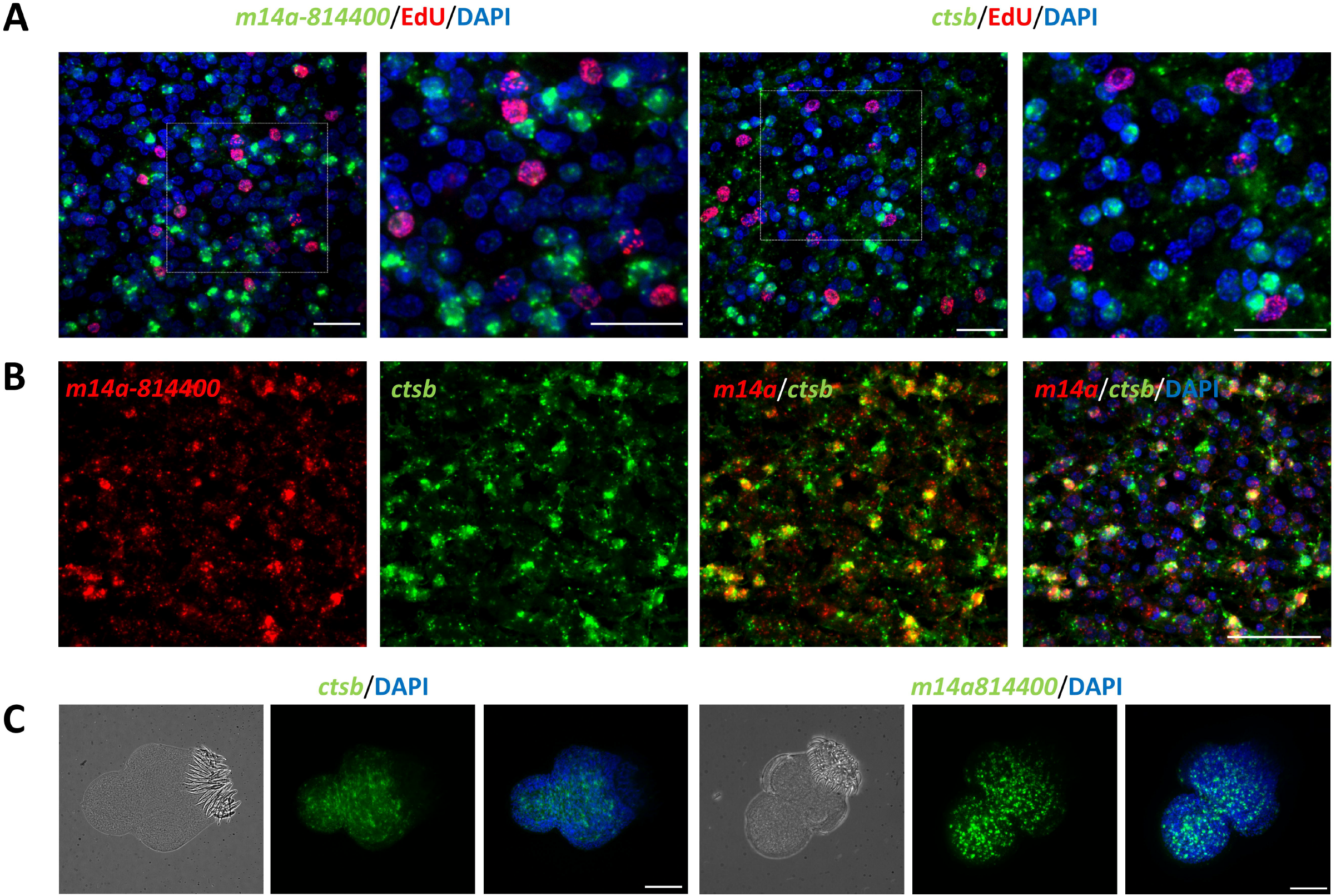
Experimental validation of *ctsb* and *m14a-814400* as molecular markers associated with putative storage cells. (A) Whole-mount *in situ* hybridization showing *m14a-814400* (green, left) and *ctsb* (green, right) expression in metacestode vesicles combined with EdU labeling (red). Parasites were exposed to EdU for 5 h prior to fixation. Nuclei were counterstained with DAPI (blue). Dashed boxes are magnified in the images to the right. Scale bars, 20 µm. (B) Double fluorescence whole-mount *in situ* hybridization showing co-expression of *m14a-814400* (red) and *ctsb* (green) in metacestode vesicles. Merged images are shown with and without DAPI nuclear staining (blue). Scale bar, 40 µm. (C) Whole-mount *in situ* hybridization showing *ctsb* (left) and *m14a-814400* (right) expression in activated protoscoleces. Bright-field, fluorescence, and merged images with DAPI nuclear staining are shown. Scale bars, 40 µm.

Together, these findings identify *ctsb* and *m14a-814400* as molecular markers associated with the putative storage cell compartment defined by scRNA-seq and provide a first spatial characterization of these transcriptionally identified populations. Although the transcriptomic atlas resolved six distinct populations within this compartment, additional molecular markers and independent functional or histological validation will be required to establish their cellular identity and determine the biological significance of their transcriptional heterogeneity (Table S10).

### Characterization of previously unassigned cell populations

In addition to the annotated cell types, the transcriptomic atlas identified two transcriptionally distinct cell populations that could not be assigned to any previously described cell type (Fig. 1B, Fig. 1C). Differential gene expression analysis revealed distinct molecular signatures for each population (Table S11).

To investigate the identity of one of these populations, we selected an annexin gene (EmuJ_000237700), one of its most highly enriched genes, for experimental validation. Whole-mount ISH revealed a striking pattern of ring-shaped structures surrounding an unlabeled central lumen and closely associated with one or more nuclei throughout the germinal layer of metacestode vesicles (Fig. 7A, Fig. 7B).

**Fig. 7.**
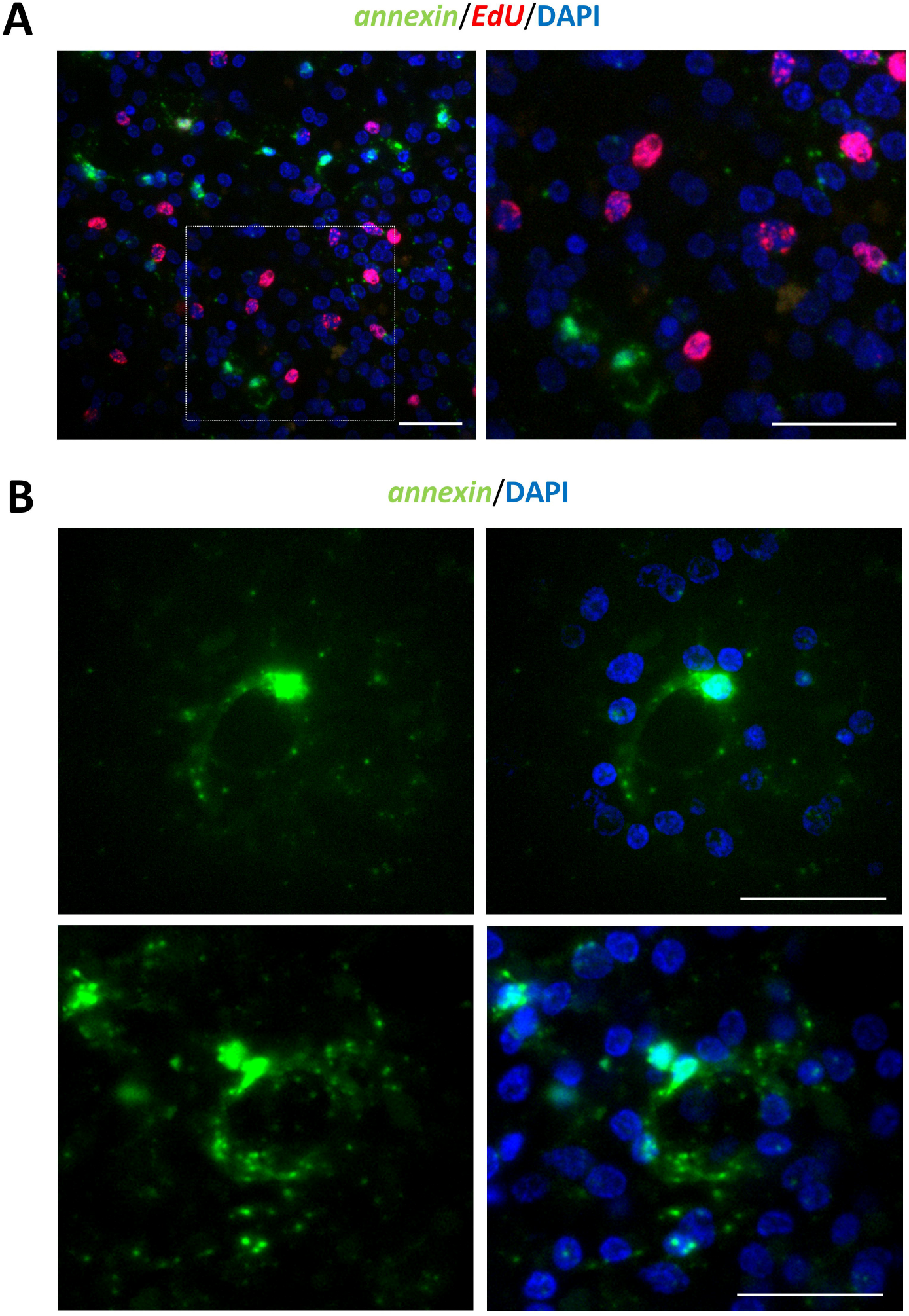
Expression pattern of *annexin* in a previously unassigned cell population. (A) Whole-mount *in situ* hybridization showing *annexin* (green) expression in metacestode vesicles combined with EdU labeling (red). Parasites were exposed to EdU for 5 h prior to fixation. Nuclei were counterstained with DAPI (blue). Dashed boxes are magnified in the images to the right. Scale bars, 20 µm. (B) Representative *annexin*-positive structure showing the characteristic ring-shaped morphology surrounding an unlabeled central lumen. Nuclei were counterstained with DAPI (blue). Scale bar, 20 µm.

The morphology of these structures closely resembled protonephridial ducts previously visualized by *Annexin* immunolocalization in Taenia crassiceps (Rios-Valencia et al., 2022). Although this morphological similarity does not establish their identity, it suggests that this previously unassigned population may correspond to cells associated with the protonephridial ducts.

## Discussion

Here, we provide the first single-cell transcriptomic atlas of the *E. multilocularis* metacestode, resolving the cellular and molecular organization of the parasite stage responsible for the persistent growth of AE lesions in the human host. The atlas resolved 26 transcriptionally distinct populations and captured the major cell types previously described at this developmental stage, including germinative, tegumental, muscle, neuronal and putative storage cells (Brehm and Koziol, 2017), together with intermediate differentiation states and previously uncharacterized populations. By combining single-cell transcriptomics with spatial validation using ISH, we further linked transcriptionally defined populations to their anatomical distribution in metacestodes and protoscoleces and established molecular markers for several major cell compartments. Together, these data reveal substantial cellular heterogeneity within the metacestode germinal layer and provide a cell-resolved framework for investigating its growth, differentiation and tissue organization.

Germinative cells are the only proliferative somatic cells of the *E. multilocularis* metacestode and underpin its continuous growth, tissue renewal and developmental plasticity (Koziol et al., 2014). The identification of *lbr*, h2a and impα as broadly expressed germinative-cell markers substantially expands the limited set of molecular tools available to study this compartment. Consistent with their association with germinative cells, all three genes were independently found to be downregulated following pharmacological depletion of proliferating cells (Herz et al., 2024). Particularly interesting is *lbr*, given the conserved role of the Lamin B receptor in peripheral heterochromatin organization and the extensive reorganization of nuclear architecture that accompanies differentiation in other metazoans (Solovei et al., 2013). Although its functional significance in *E. multilocularis* remains unknown, its association with germinative cells offers a molecular entry point for investigating changes in nuclear organization during parasite cell differentiation.

Beyond the identification of new markers, our atlas resolves the molecular heterogeneity previously observed within this morphologically homogeneous compartment (Koziol et al., 2014; Herz et al., 2024). Notably, genes previously associated with restricted subsets of germinative cells, including nos1, kal1, post2, npp27 and math6, were preferentially associated with different stem-cell populations. This organization suggests that the previously recognized heterogeneity reflects distinct biological states within the germinative-cell compartment. Particularly informative is Stem2, in which nos1 and kal1 converge within the same transcriptional state. Given the previous association of kal1 with slowly cycling germinative cells (Herz et al., 2024), the transcriptional profile of Stem2 is consistent with a relatively undifferentiated and potentially slow-cycling germinative-cell state. Its position within the developmental hierarchy, however, remains to be established. Stem3 and Stem4 point to a different aspect of germinative-cell heterogeneity, as both showed transcriptional biases toward lineage-associated programs—tegumental and muscle, respectively. This resembles the lineage-biased heterogeneity of planarian neoblasts, in which subsets of proliferative cells express transcriptional programs associated with particular differentiated cell types (Scimone et al., 2014; Fincher et al., 2018). Such biases may reflect early stages of differentiation, but do not by themselves establish lineage commitment or developmental potential (Raz et al., 2021). By contrast, the biological significance of the distinct transcriptional programs observed in Stem1 and Stem5 remains to be resolved.

The differential representation of Stem3 between parasite isolates adds a developmental dimension to germinative-cell heterogeneity. Stem3 was readily resolved in LS23, which retains the capacity to form brood capsules and protoscoleces, but was nearly absent from the developmentally deficient H95 isolate. Subclustering of LS23 further identified a Stem3-like state enriched in npp27, previously detected in a subset of germinative cells (Herz et al., 2024). Together with the lineage-associated profile of Stem3, these observations raise the possibility that the representation of particular germinative-cell states is linked to developmental competence. However, whether the differential representation of Stem3 reflects genuine differences in the germinative-cell composition of the two isolates, and how these relate to their developmental capacity, remains to be established.

The transcriptional profiles of differentiated cell populations also provide insight into functions that are not readily apparent from morphology alone. This is particularly evident for muscle. The presence of muscle fibers in metacestode vesicles has been enigmatic, since the fibers are disorganized and the vesicles lack motility (Koziol et al., 2013). However, previous studies showed that *E. multilocularis* muscle cells are a source of Wnt ligands involved in anterior-posterior patterning and suggested that this signaling function may account for the retention of muscle in the otherwise immobile metacestode (Koziol et al., 2016a). Consistent with this hypothesis, Wnt signaling was among the biological processes significantly enriched in the muscle populations in our dataset. Components of the TGF-β pathway and homologues of Hedgehog ligands were also preferentially expressed in these populations, suggesting that the signaling role of muscle may extend beyond the Wnt pathway. This organization parallels that of planarians, where muscle cells express position-control genes from multiple signaling pathways and provide positional information required for tissue patterning and regeneration (Witchley et al., 2013; Scimone et al., 2016; Fincher et al., 2018), and is consistent with the association of developmental signaling pathways with muscle populations in S. mansoni (Wendt et al., 2020). Recent functional evidence that canonical WNT signaling is essential for *E. multilocularis* metacestode development further supports the relevance of this signaling system to larval patterning (Herrmann et al., 2026). Together, these findings strengthen the view of metacestode muscle as an important source of developmental signals and suggest that this role extends beyond Wnt signaling.

Both muscle populations also displayed a prominent extracellular-matrix transcriptional signature. In planarians, muscle is a major source of ECM components and functions as a connective tissue that contributes to the maintenance of tissue organization (Cote et al., 2019). ECM-associated transcriptional programs have likewise been detected in muscle populations of S. mansoni (Wendt et al., 2020). The coexistence of developmental signaling and ECM programs therefore points to a broader contribution of *E. multilocularis* muscle to tissue organization. The transcriptional differences between the two muscle populations further suggest specialization within this compartment, although their spatial and functional significance remains to be established.

The putative neuronal population identified in our atlas provides molecular access to a particularly poorly understood component of the metacestode. Previous anatomical studies revealed a simple subtegumental nerve net in the germinal layer, a striking feature of this largely immotile larval stage whose function remains unclear and for which a possible neuroendocrine role has been proposed (Koziol et al., 2013). The expression of *nrx1α* and *meis1*, together with the neurite-like morphology of *nrx1α*-positive cells and the association of these markers with the nervous system in protoscoleces, supports the neuronal assignment of this population. This interpretation is further supported by the conserved role of neurexins as presynaptic cell-adhesion molecules in bilaterians (Guzman et al., 2024) and by the established roles of Meis-family transcription factors in nervous-system development and neuronal specification (Campbell and Walthall, 2016). Whereas both genes showed largely overlapping expression in the metacestode, their patterns diverged markedly during protoscolex development. *nrx1α* broadly labelled the protoscolex nervous system, while *meis1* was restricted to cells surrounding the rostellum. This divergence is particularly interesting because the protoscolex nervous system develops independently from the metacestode nerve net and acquires a substantially more complex organization (Koziol et al., 2013). Previous analyses have likewise revealed distinct spatial expression of multiple neuropeptide genes within the protoscolex nervous system (Koziol et al., 2016b). Together, these observations suggest that the development of the protoscolex nervous system involves the emergence of molecularly distinct neuronal domains. The markers identified here provide new tools with which this organization can now be investigated.

The tegument represents another compartment in which the atlas revealed both developmental and transcriptional heterogeneity. Previous studies identified *muc1* and alp-2 as markers of differentiated tegumental cells in the metacestode (Koziol et al., 2014). The identification of *tsp-355900* and *teg-372400* expands this repertoire and reveals marked changes in tegumental gene expression during larval development. In particular, the persistence of *teg-372400* and loss of *tsp-355900* expression in protoscoleces, together with the previously reported absence of *muc1* and alp-2 from this stage (Koziol et al., 2014), indicate substantial molecular remodeling of the tegument during the transition from metacestode to protoscolex.

The tegumental populations also shared a prominent microtubule-associated program, including dynein light-chain and tubulin genes. Similar transcriptional signatures have been recovered from tegumental populations across several developmental stages of S. mansoni (Wendt et al., 2020; Diaz Soria et al., 2020; Diaz Soria et al., 2024; Attenborough et al., 2024), suggesting that microtubule-associated functions may represent a conserved component of the tegumental program in parasitic flatworms. Despite this shared program, the resolution of three tegumental populations indicates molecular heterogeneity among the cytons underlying the syncytial tegument. This is consistent with recent comparative analyses of cestodes showing regionally restricted gene expression among tegumental cytons and suggesting functional specialization within an otherwise continuous syncytium (Guarnaschelli et al., 2026). Notably, Tegument 1 preferentially expressed several membrane transporters, including the SLC5-family transporter EmuJ_000714000. This gene and the more broadly expressed concentrative nucleoside transporter EmuJ_000127200 are orthologous to candidate tegument-associated nutrient transporters identified in other cestodes and are strongly upregulated following protoscolex activation and culture (Guarnaschelli et al., 2026). The enrichment of transporters in Tegument 1 is consistent with the central role of the cestode tegument in nutrient uptake and the extensive metabolic dependence of tapeworms on host-derived nutrients (Tsai et al., 2013). In contrast, Tegument 3 preferentially expressed several dynein light-chain genes. Together, these differences raise the possibility of functional specialization among tegumental cytons, although the physiological roles of the individual populations remain to be established.

The localized *teg-372400*^high^/*muc1*^low^ domains observed within the germinal layer may represent a spatial correlate of tegumental heterogeneity and/or developmental remodeling. Their relationship to brood-capsule development is particularly intriguing because *muc1* expression is lost from early brood-capsule buds and developing protoscoleces (Koziol et al., 2014), whereas *teg-372400* expression is maintained during this developmental transition. Although these domains lack the morphological characteristics of recognizable brood capsules, the local divergence between two otherwise overlapping tegumental markers raises the possibility that molecular changes in tegumental identity precede overt morphological differentiation. Whether these domains represent an early stage of developmental remodeling will require direct analysis of their fate over time.

The putative storage-cell populations present a different challenge, because their assignment currently rests primarily on their transcriptional profile rather than on direct correspondence with the morphologically defined storage cells of the germinal layer. Nevertheless, their prominent metabolic signature is consistent with the presumed nutrient-storage function of these cells (Koziol et al., 2014). Particularly noteworthy is the expression of genes involved in lipid handling, including fatty acid-binding proteins and Antigen B family members, given the restricted lipid biosynthetic capacity of Echinococcus and its dependence on host-derived lipids (Tsai et al., 2013). Both protein families have established lipid-binding and transport properties in Echinococcus (Alvite and Esteves, 2012; Obal et al., 2012; Silva-Álvarez et al., 2015; Alvite and Esteves, 2016). Together with the enrichment of glycolytic and glucose metabolic processes, these features are consistent with a metabolically specialized compartment involved in nutrient processing and storage.

A similarly prominent metabolic signature has been reported for parenchymal populations in S. mansoni, where genes associated with carbohydrate and lipid metabolism accompany the presence of carbohydrate particles and lipid droplets (Diaz Soria et al., 2024). Cathepsin B-like genes have likewise been associated with parenchymal populations across S. mansoni developmental stages (Diaz Soria et al., 2020; Wendt et al., 2020; Attenborough et al., 2024), providing additional comparative context for the association of *ctsb* with the putative storage compartment in our atlas. The six populations identified here further point to substantial transcriptional heterogeneity within this putative compartment, although the biological basis of this diversity remains unclear. Direct association of *ctsb* and *m14a-814400* expression with the characteristic lipid and/or glycogen stores of morphologically defined storage cells would provide stronger evidence for their identity.

Finally, the atlas provides molecular access to populations that could not be assigned using previously available markers. Particularly striking was the distribution of the *annexin* transcript associated with one of these populations, which delineated ring-like structures resembling components of the protonephridial system. Although this system has been described morphologically in *E. multilocularis*, the molecular identity of cells associated with the protonephridial ducts remains poorly characterized (Koziol et al., 2013). The localization of *Annexin* to protonephridial ducts and flame cells in T. crassiceps provides additional support for this interpretation (Rios-Valencia et al., 2022). These observations raise the possibility that the atlas has captured a molecular signature of cells associated with the protonephridial ducts, opening this poorly characterized compartment to further molecular investigation.

Overall, this study moves our understanding of the *E. multilocularis* metacestode beyond a largely morphology-based description toward a cell-resolved view of its molecular organization. The heterogeneity resolved within germinative and differentiated compartments reveals an organization that is not apparent from morphology alone, while the markers identified here make several of these populations experimentally accessible. In particular, distinct germinative-cell states provide a basis for dissecting the organization of proliferation and differentiation, while the specialized transcriptional programs of differentiated populations point to diverse roles in signaling, tissue organization and metabolism. Together, these findings establish a cellular basis for investigating the mechanisms that sustain the remarkable growth and developmental plasticity of the *E. multilocularis* metacestode within the host.

## Acknowledgements

The authors thank Dirk Radloff for excellent technical assistance and Dr. Fabian Imdahl (Helmholtz Institute for RNA-based Infection Research) for valuable advice and assistance with cell sorting.

## Supplementary Video

Video S1. Three-dimensional visualization of *nrx1α* expression (green) in an activated protoscolex.

---

Fig. S1. Gene Ontology (GO) Biological Process enrichment analysis of the major cell populations identified in the metacestode cell atlas. Heatmap showing representative enriched GO Biological Process terms associated with the major cell populations identified by single-cell RNA sequencing. Columns correspond to the major annotated cell populations (Tegument, Storage, Stem, Progeny, Intermediate, Neuron-like, Muscle, and Unknown), and rows represent selected enriched GO terms grouped according to the population in which they showed the strongest enrichment. Color intensity indicates the scaled enrichment score for each GO term across cell populations.

Fig. S2. Experimental validation of additional germinative cell markers. (A) Whole-mount *in situ* hybridization showing *impα* (green, left) and *h2a* (green, right) expression in metacestode vesicles combined with EdU labeling (red). Parasites were exposed to EdU for 5 h prior to fixation. Nuclei were counterstained with DAPI (blue). Dashed boxes are magnified in the images to the right. Scale bars, 20 µm. (B) Double fluorescence whole-mount *in situ* hybridization showing co-expression of *impα* (red) and the germinative cell marker *cip2ah* (green) in metacestode vesicles. Merged images are shown with and without DAPI nuclear staining (blue). Scale bar, 40 µm. (C) Double fluorescence whole-mount *in situ* hybridization showing co-expression of *h2a* (red) and the germinative cell marker *cip2ah* (green) in metacestode vesicles. Merged images are shown with and without DAPI nuclear staining (blue). Scale bar, 40 µm.

Fig. S3. Experimental validation of *nkx2.5* expression in muscle cells. (A) Whole-mount *in situ* hybridization showing *nkx2.5* expression (green) in metacestode vesicles combined with EdU labeling (red). Parasites were exposed to EdU for 5 h prior to fixation. Nuclei were counterstained with DAPI (blue). Dashed boxes are magnified in the images to the right. Scale bars, 20 µm. (B) Whole-mount *in situ* hybridization showing the distribution of *nkx2.5* expression in an activated protoscolex. Bright-field, fluorescence, and merged images with DAPI nuclear staining are shown. Scale bar, 40 µm. (C) Combined detection of *nkx2.5* transcripts by whole-mount *in situ* hybridization (green) and Tropomyosin immunofluorescence (red) in metacestode vesicles. Merged images are shown with and without DAPI nuclear staining (blue). Scale bar, 20 µm. (D) Double fluorescence whole-mount *in situ* hybridization showing overlapping expression of *nkx2.5* (red) and titin (green) in metacestode vesicles. Cells with strong expression of both markers, as well as cells displaying comparatively weaker signal for either *nkx2.5* or titin, are visible. The merged image is shown with DAPI nuclear staining (blue). Scale bar, 40 µm.

Fig. S4. Expression of the tegumental markers *tsp-355900* and *teg-372400* during larval development. (A, B) Whole-mount *in situ* hybridization showing *tsp-355900* expression (green) in metacestode vesicles containing brood capsules and protoscoleces. *tsp-355900* expression was detected in the metacestode germinal layer and brood-capsule wall but was not detected in protoscoleces. (C, D) Whole-mount *in situ* hybridization showing *teg-372400* expression (green) in developing brood capsules. Nuclei were counterstained with DAPI (blue), and merged images are shown in the right panels. Scale bars, 40 µm.

## Supplementary Tables

Table S1. Oligonucleotide primer sequences used for WISH.

Table S2. List of samples used for single-cell RNA sequencing.

Table S3. Marker genes for each scRNA-seq cluster identified using Seurat FindAllMarkers with the Wilcoxon rank-sum test.

Table S4. Marker genes for each scRNA-seq cluster identified using Seurat FindAllMarkers with receiver operating characteristic (ROC) analysis, including area under the curve (AUC) scores.

Table S5. Marker genes distinguishing the Stem cell scRNA-seq clusters identified using Seurat FindAllMarkers with the Wilcoxon rank-sum test.

Table S6. Marker genes for H95 stem cell subclusters identified by isolate-specific subclustering using Seurat FindAllMarkers with the Wilcoxon rank-sum test.

Table S7. Marker genes for LS23 stem cell subclusters identified by isolate-specific subclustering using Seurat FindAllMarkers with the Wilcoxon rank-sum test.

Table S8. Marker genes distinguishing the Muscle 1 and Muscle 2 scRNA-seq clusters identified using Seurat FindAllMarkers with the Wilcoxon rank-sum test.

Table S9. Marker genes distinguishing the Tegument 1, Tegument 2, and Tegument 3 scRNA-seq clusters identified using Seurat FindAllMarkers with the Wilcoxon rank-sum test.

Table S10. Marker genes distinguishing the Storage 1-6 scRNA-seq clusters identified using Seurat FindAllMarkers with the Wilcoxon rank-sum test.

Table S11. Marker genes distinguishing the Unknown 1 and Unknown 2 scRNA-seq clusters identified using Seurat FindAllMarkers with the Wilcoxon rank-sum test.

## References

Alvite, G., & Esteves, A. (2012). Lipid binding proteins from parasitic platyhelminthes. Frontiers in Physiology, 3, 363.

Alvite, G., & Esteves, A. (2016). Echinococcus granulosus fatty acid binding proteins subcellular localization. Experimental Parasitology, 164, 1–4.

Attenborough, T., Rawlinson, K. A., Diaz Soria, C. L., Ambridge, K., Sankaranarayanan, G., Graham, J., et al. (2024). A single-cell atlas of the miracidium larva of Schistosoma mansoni reveals cell types, developmental pathways, and tissue architecture. eLife, 13, RP95628.

Brehm, K., & Koziol, U. (2014). On the importance of targeting parasite stem cells in anti-echinococcosis drug development. Parasite, 21, 72.

Brehm, K., & Koziol, U. (2017). Echinococcus-host interactions at cellular and molecular levels. Advances in Parasitology, 95, 147–212.

Brunetti, E., Kern, P., & Vuitton, D. A. (2010). Expert consensus for the diagnosis and treatment of cystic and alveolar echinococcosis in humans. Acta Tropica, 114, 1–16.

Campbell, R. F., & Walthall, W. W. (2016). Meis/UNC-62 isoform dependent regulation of CoupTF-II/UNC-55 and GABAergic motor neuron subtype differentiation. Developmental Biology, 419(2), 250–261.

Casulli, A., Barth, T. F., & Tamarozzi, F. (2019). Echinococcus multilocularis. Trends in Parasitology, 35(9), 738–739.

Cote, L. E., Simental, E., & Reddien, P. W. (2019). Muscle functions as a connective tissue and source of extracellular matrix in planarians. Nature Communications, 10, 1592.

Deplazes, P., Rinaldi, L., Alvarez Rojas, C. A., Torgerson, P. R., Harandi, M. F., Romig, T., et al. (2017). Global distribution of alveolar and cystic echinococcosis. Advances in Parasitology, 95, 315–493.

Diaz Soria, C. L., Lee, J., Chong, T., Coghlan, A., Tracey, A., Young, M. D., et al. (2020). Single-cell atlas of the first intra-mammalian developmental stage of the human parasite Schistosoma mansoni. Nature Communications, 11, 6411.

Diaz Soria, C. L., Attenborough, T., Lu, Z., Fontenla, S., Graham, J., Hall, C., et al. (2024). Single-cell transcriptomics of the human parasite Schistosoma mansoni first intra-molluscan stage reveals tentative tegumental and stem-cell regulators. Scientific Reports, 14, 5974.

Fincher, C. T., Wurtzel, O., de Hoog, T., Kravarik, K. M., & Reddien, P. W. (2018). Cell type transcriptome atlas for the planarian Schmidtea mediterranea. Science, 360, eaaq1736.

Guarnaschelli, I., Lima, A., Velazco, R., Bergmann, M., Preza, M., Calvelo, J., et al. (2026). Comparative proteomics reveals a conserved core of tegumental proteins in parasitic flatworms. bioRxiv.

Guzman, C., Mohri, K., Nakamura, R., Miyake, M., Tsuchiya, Y., Tomii, K., & Watanabe, H. (2024). Neuronal and non-neuronal functions of the synaptic cell adhesion molecule neurexin in Nematostella vectensis. Nature Communications, 15, 6495.

Hemphill, A., Stadelmann, B., Rufener, R., Spiliotis, M., Boubaker, G., Müller, J., et al. (2014). Treatment of echinococcosis: albendazole and mebendazole—what else? Parasite, 21, 70.

Herrmann, R., Herz, M., Rudolf, K., Koike, A., Spiliotis, M., Bergmann, M., et al. (2026). Canonical WNT signalling governs Echinococcus metacestode development. PLOS Pathogens, 22(3), e1014046.

Herz, M., Zarowiecki, M., Wessels, L., Pätzel, K., Herrmann, R., Braun, C., et al. (2024). Genome-wide transcriptome analysis of *Echinococcus multilocularis* larvae and germinative cell cultures reveals genes involved in parasite stem cell function. Frontiers in Cellular and Infection Microbiology, 14, 1335946.

Kern, P., Menezes da Silva, A., Akhan, O., Müllhaupt, B., Vizcaychipi, K. A., Budke, C., et al. (2017). The echinococcoses: diagnosis, clinical management and burden of disease. Advances in Parasitology, 96, 259–369.

Koziol, U., & Brehm, K. (2015). Recent advances in Echinococcus genomics and stem cell research. Veterinary Parasitology, 213(3–4), 92–102.

Koziol, U., Jarero, F., Olson, P. D., & Brehm, K. (2016a). Comparative analysis of Wnt expression identifies a highly conserved developmental transition in flatworms. BMC Biology, 14, 10.

Koziol, U., Koziol, M., Preza, M., Costábile, A., Brehm, K., & Castillo, E. (2016b). De novo discovery of neuropeptides in the genomes of parasitic flatworms using a novel comparative approach. International Journal for Parasitology, 46(11), 709–721.

Koziol, U., Krohne, G., & Brehm, K. (2013). Anatomy and development of the larval nervous system in *Echinococcus multilocularis*. Frontiers in Zoology, 10, 24.

Koziol, U., Radio, S., Smircich, P., Zarowiecki, M., Fernández, C., & Brehm, K. (2015). A novel terminal-repeat retrotransposon in miniature (TRIM) is massively expressed in *Echinococcus multilocularis* stem cells. Genome Biology and Evolution, 7(8), 2136–2153.

Koziol, U., Rauschendorfer, T., Zanon Rodríguez, L., Krohne, G., & Brehm, K. (2014). The unique stem cell system of the immortal larva of the human parasite *Echinococcus multilocularis*. EvoDevo, 5, 10.

Lähnemann, D., Köster, J., Szczurek, E., McCarthy, D. J., Hicks, S. C., Robinson, M. D., et al. (2020). Eleven grand challenges in single-cell data science. Genome Biology, 21, 31.

Lascano, E. F., Coltorti, E. A., & Varela-Diaz, V. M. (1975). Fine structure of the germinal membrane of Echinococcus granulosus cysts. Journal of Parasitology, 61, 853–860.

Li, P., Nanes Sarfati, D., Xue, Y., Yu, X., Tarashansky, A. J., Quake, S. R., & Wang, B. (2021). Single-cell analysis of Schistosoma mansoni identifies a conserved genetic program controlling germline stem cell fate. Nature Communications, 12, 485.

Molinaro, A. M., & Pearson, B. J. (2016). In silico lineage tracing through single cell transcriptomics identifies a neural stem cell population in planarians. Genome Biology, 17, 87.

Obal, G., Ramos, A. L., Silva, V., Lima, A., Batthyány, C., Bessio, M. I., et al. (2012). Characterisation of the native lipid moiety of Echinococcus granulosus antigen B. PLoS Neglected Tropical Diseases, 6, e1642.

Plass, M., Solana, J., Wolf, F. A., Ayoub, S., Misios, A., Glažar, P., et al. (2018). Cell type atlas and lineage tree of a whole complex animal by single-cell transcriptomics. Science, 360, eaaq1723.

Puckelwaldt, O., Gramberg, S., Ajmera, S., Koepke, J., Shamsara, J., Samakovlis, C., et al. (2026). Single-cell transcriptomics identifies a p21-activated kinase important for survival of the zoonotic parasite Fasciola hepatica. iScience, 29, 116778.

Raz, A. A., Wurtzel, O., & Reddien, P. W. (2021). Planarian stem cells specify fate yet retain potency during the cell cycle. Cell Stem Cell, 28, 1307–1322.

Rios-Valencia, D. G., Mompala-García, Y., Marquez-Navarro, A., Tirado-Mendoza, R., & Ambrosio, J. (2022). *Annexin* in Taenia crassiceps ORF strain is localized in the osmoregulatory system. Acta Parasitologica, 67, 827–834.

Sakamoto, T., & Sugimura, M. (1970). Studies on echinococcosis XXIII. Electron microscopical observations on histogenesis of larval *Echinococcus multilocularis*. Japanese Journal of Veterinary Research, 18, 131–144.

Schubert, A., Koziol, U., Cailliau, K., Vanderstraete, M., Dissous, C., & Brehm, K. (2014). Targeting *Echinococcus multilocularis* stem cells by inhibition of the Polo-like kinase EmPlk1. PLoS Neglected Tropical Diseases, 8, e2870.

Scimone, M. L., Cote, L. E., Rogers, T., & Reddien, P. W. (2016). Two FGFRL-Wnt circuits organize the planarian anteroposterior axis. eLife, 5, e12845.

Scimone, M. L., Kravarik, K. M., Lapan, S. W., & Reddien, P. W. (2014). Neoblast specialization in regeneration of the planarian Schmidtea mediterranea. Stem Cell Reports, 3, 339–352.

Silva-Álvarez, V., Folle, A. M., Ramos, A. L., Zamarreño, F., Costabel, M. D., García-Zepeda, E., et al. (2015). Echinococcus granulosus antigen B: A hydrophobic ligand binding protein at the host–parasite interface. Prostaglandins, Leukotrienes and Essential Fatty Acids, 93, 17–23.

Solovei, I., Wang, A. S., Thanisch, K., Schmidt, C. S., Krebs, S., Zwerger, M., et al. (2013). LBR and lamin A/C sequentially tether peripheral heterochromatin and inversely regulate differentiation. Cell, 152, 584–598.

Spiliotis, M., & Brehm, K. (2009). Axenic *in vitro* cultivation of *Echinococcus multilocularis* metacestode vesicles and the generation of primary cell cultures. Methods in Molecular Biology, 470, 245–262.

Stuart, T., & Satija, R. (2019). Integrative single-cell analysis. Nature Reviews Genetics, 20, 257–272.

Swapna, L. S., Molinaro, A. M., Lindsay-Mosher, N., Pearson, B. J., & Parkinson, J. (2018). Comparative transcriptomic analyses and single-cell RNA sequencing of the freshwater planarian Schmidtea mediterranea identify major cell types and pathway conservation. Genome Biology, 19, 124.

Thompson, R. C. A. (2017). Biology and systematics of Echinococcus. Advances in Parasitology, 95, 65–109.

Tsai, I. J., Zarowiecki, M., Holroyd, N., Garciarrubio, A., Sánchez-Flores, A., Brooks, K. L., et al. (2013). The genomes of four tapeworm species reveal adaptations to parasitism. Nature, 496, 57–63.

van Wolfswinkel, J. C., Wagner, D. E., & Reddien, P. W. (2014). Single-cell analysis reveals functionally distinct classes within the planarian stem cell compartment. Cell Stem Cell, 15, 326–339.

Wang, B., Lee, J., Li, P., Saberi, A., Yang, H., Liu, C., et al. (2018). Stem cell heterogeneity drives the parasitic life cycle of Schistosoma mansoni. eLife, 7, e35449.

Wendt, G. R., Zhao, L., Chen, R., Liu, C., O’Donoghue, A. J., Caffrey, C. R., et al. (2020). A single-cell RNA-seq atlas of Schistosoma mansoni identifies a key regulator of blood feeding. Science, 369, 1644–1649.

Witchley, J. N., Mayer, M., Wagner, D. E., Owen, J. H., & Reddien, P. W. (2013). Muscle cells provide instructions for planarian regeneration. Cell Reports, 4, 633–641.

Zeng, A., Li, H., Guo, L., Gao, X., McKinney, S., Wang, Y., et al. (2018). Prospectively isolated Tetraspanin+ neoblasts are adult pluripotent stem cells underlying planaria regeneration. Cell, 173(7), 1593–1608.e20.

