## Supplementary figures and images for "A single-cell transcriptomic atlas of the Echinococcus multilocularis metacestode reveals cellular diversity and molecular specialization"

### Fig. S1

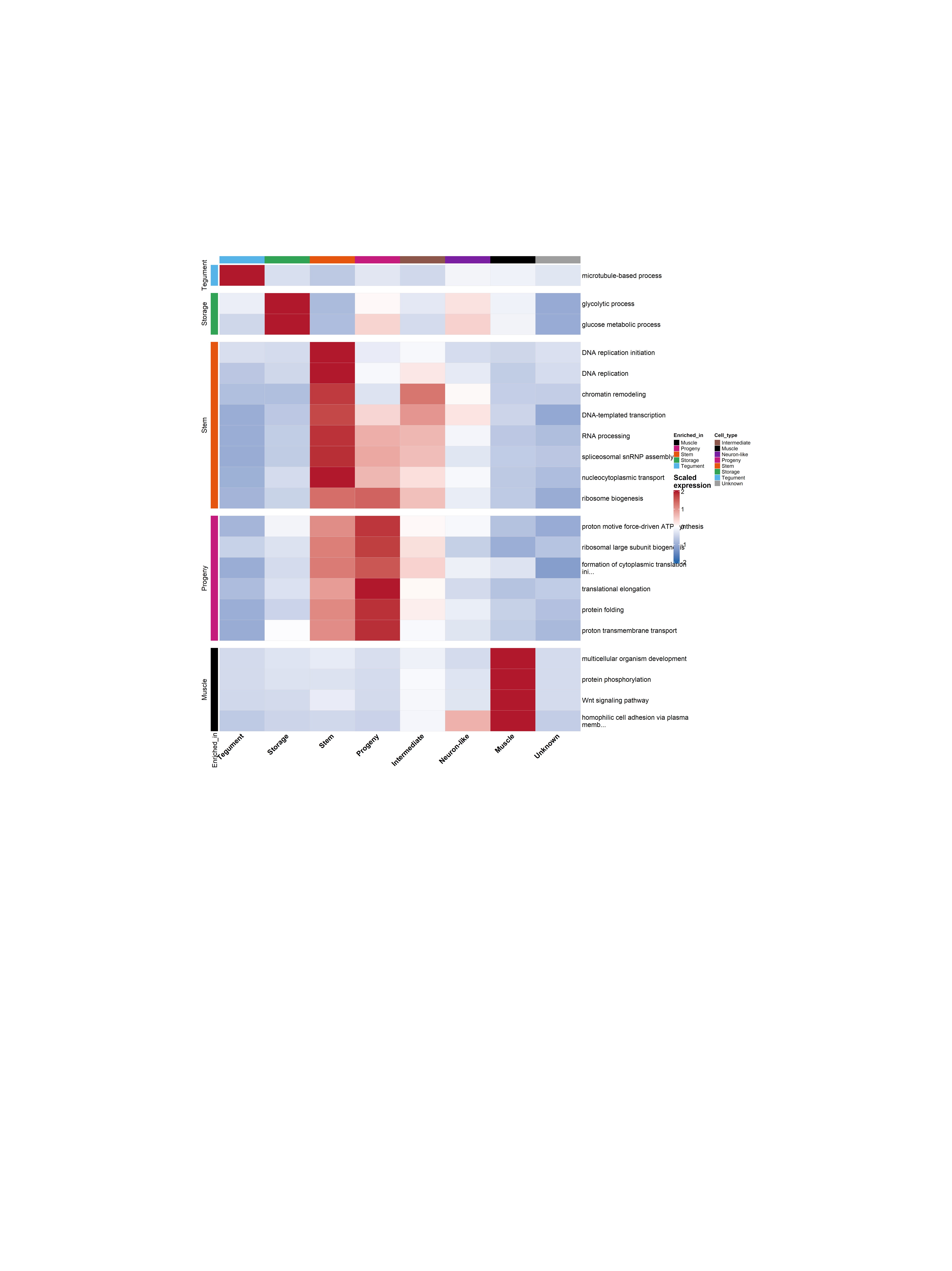

### Fig. S2

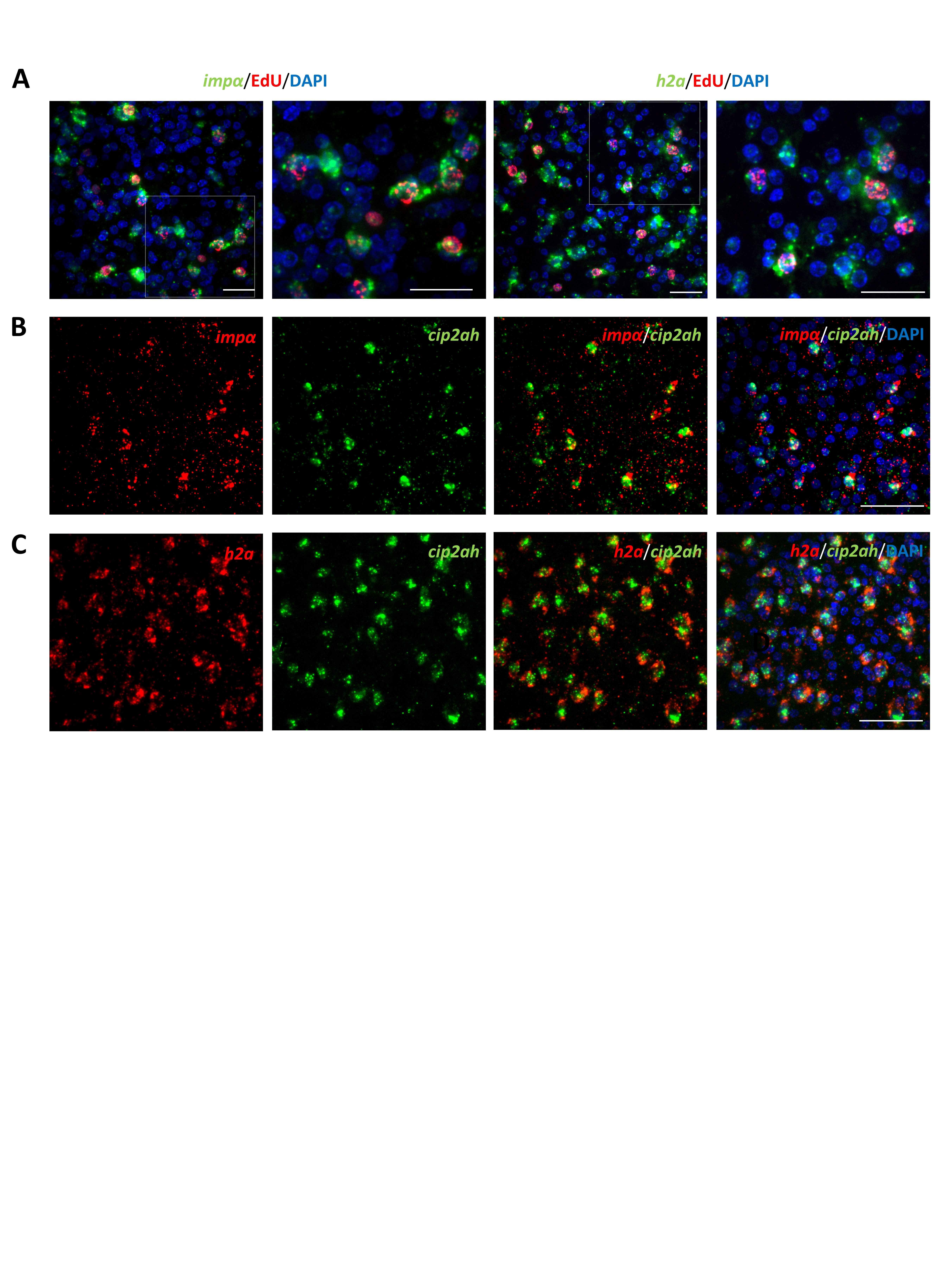

### Fig. S3

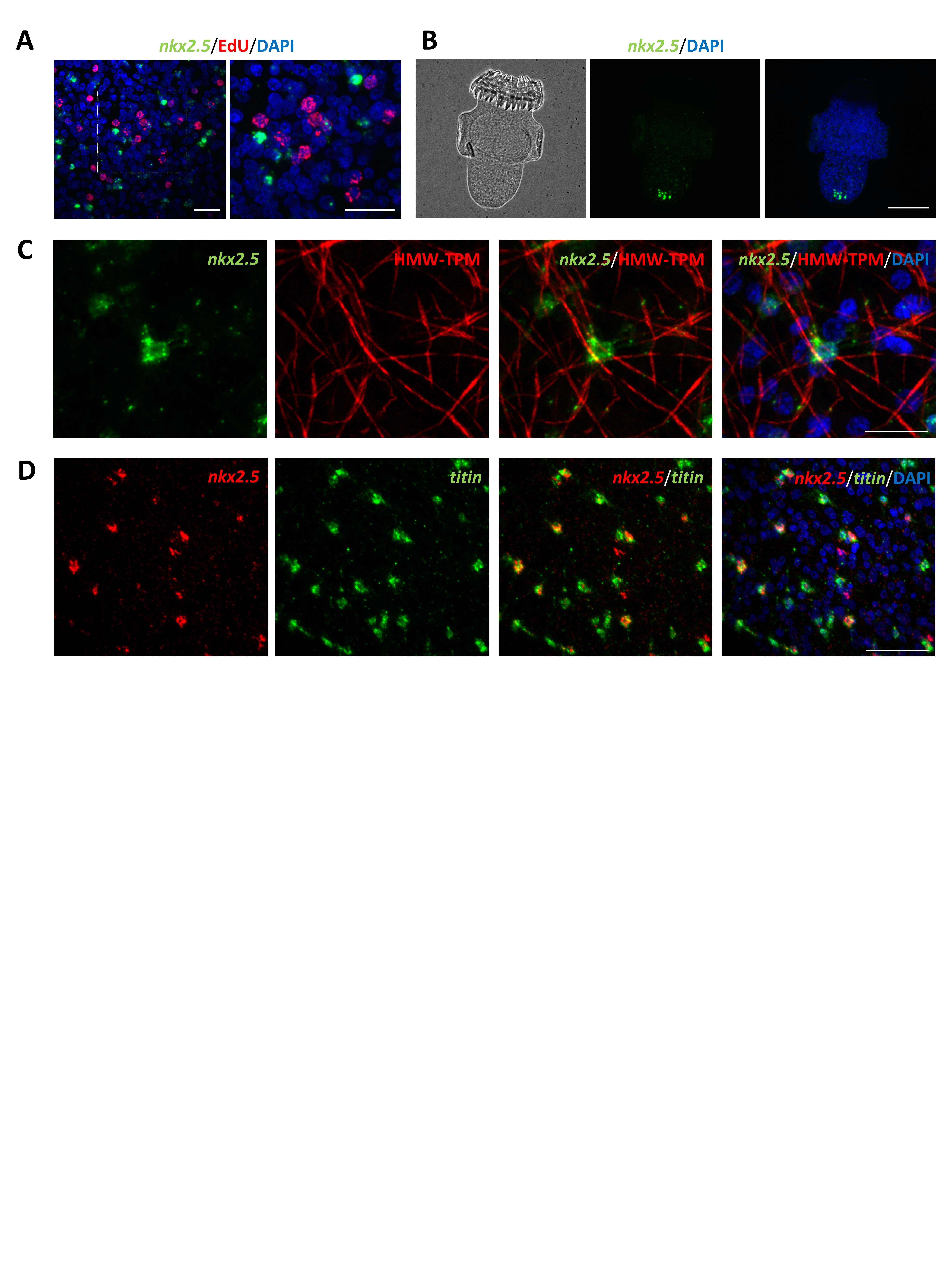

### Fig. S4

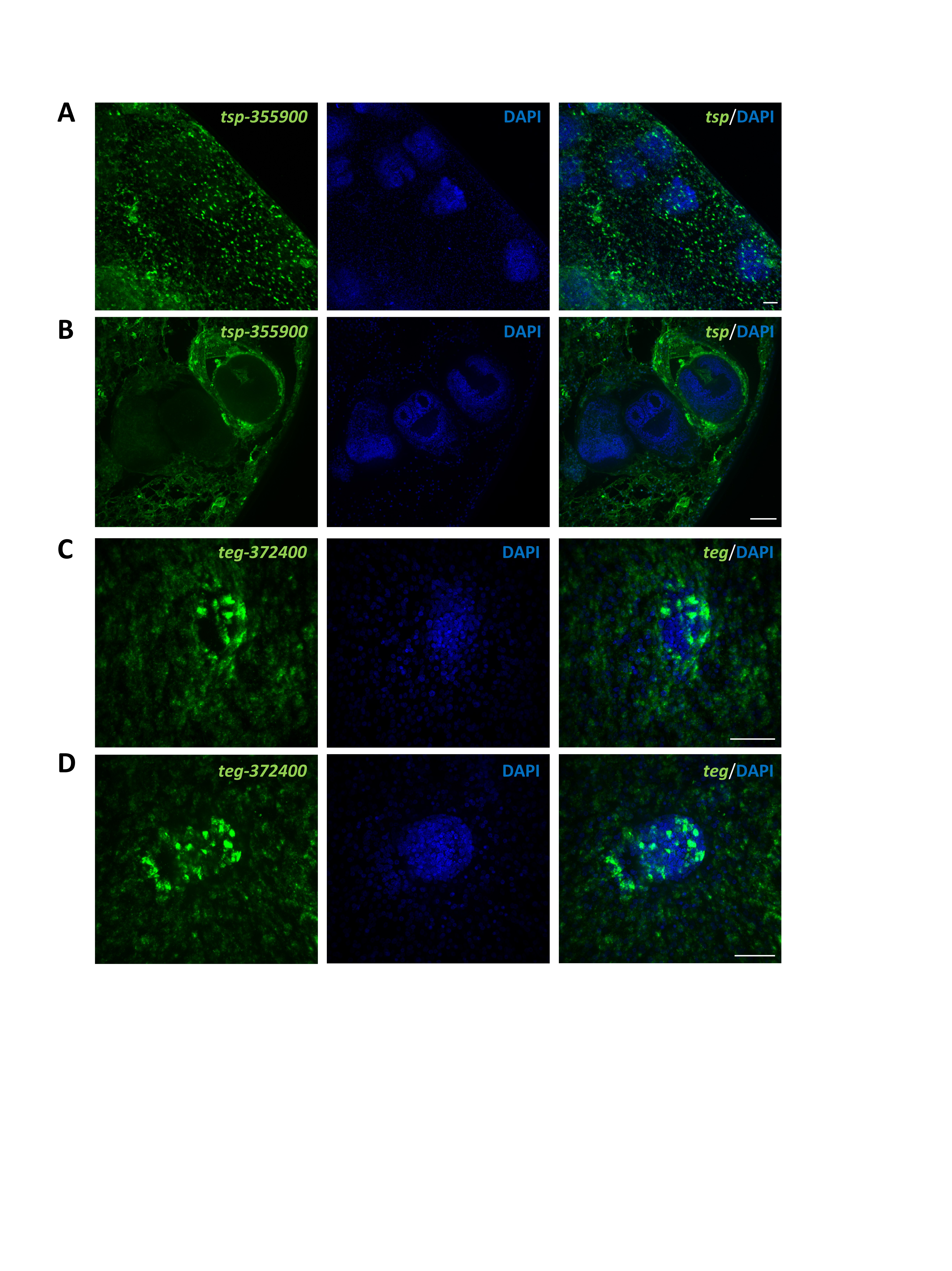
